# Plant and Aphid Genotypes Modulate Legume Rhizobium-Induced Defense Against Aphids

**DOI:** 10.64898/2025.12.22.695904

**Authors:** Gaurav Pandharikar, Hugo Mathe-Hubert, Jean-Luc Gatti, Jean-Christophe Simon, Marylène Poirié, Pierre Frendo

**Author notes:** <u>Corresponding author</u>: Pandharikar Gaurav, <u>Address:</u> Sophia Agrobiotech Institute (ISA), UCA / CNRS / INRAE, 400 route des Chappes, 06903 Sophia Antipolis, France. These authors are co-last authors (Marylène Poirié, Pierre Frendo).

## Abstract

Sustainable protein production is needed to ensure food security while mitigating climate change. Leguminous crops contribute to these goals by providing protein-rich seeds and improving soil fertility through nitrogen-fixing symbiosis (NFS) with rhizobia. Beyond their nutritional and agronomic benefits, legumes face major challenges from insect herbivores such as the pea aphid, *Acyrthosiphon pisum*. While NFS can enhance plant defences against aphids, the influence of plant and aphid genotypic variation on these rhizobia-mediated effects remains poorly understood. Here, we investigated how genotype interactions influence rhizobia-mediated defence priming by using two *Medicago truncatula* genotypes (A17 and R108) grown under NFS with *Sinorhizobium meliloti* 2011 or nitrate (NI) feeding conditions and infested with three clonal lines representing two pea aphid biotypes. We analysed aphid performance, plant fitness, and plant leaf expression of defence genes associated with the jasmonic acid (JA) and salicylic acid (SA) signalling pathways over a 12-day period following aphid infestation. Aphid fitness varied significantly with both plant and aphid genotypes and was further modulated by rhizobia inoculation. Expression of defence marker genes involved in SA and JA pathway was dependent on specific plant–aphid genotype combinations and was significantly modulated by rhizobia inoculation. This pattern suggests that both SA- and JA-mediated plant defences contribute to regulating aphid weight, possibly through different mechanisms or in response to different plant–aphid interactions. Our study shows that plant and aphid genotypes, rhizobia inoculation and interactions (G × G × R) are thus central components driving plantaphid-rhizobia interaction dynamics. Our study highlights importance of genetic context and microbial symbiosis in structuring multitrophic interactions and suggests new opportunities to optimize pest resistance using plant-beneficial microbial associations.

## 1. INTRODUCTION

Legumes are an important source of protein for both humans and animals. Their capacity to establish symbiotic associations with nitrogen-fixing microbes allows them to decrease dependence on synthetic nitrogen fertilizers, contributing to climate change mitigation. However, legumes like many other crops are attacked by different pests, among which aphids represent a major threat. Aphids reduce the crop productivity both by causing severe damage to plants through direct feeding or more critically, being vectors for many plant viral diseases. Aphid species differ in their host range, some are polyphagous, feeding on different plant species, while others are restricted to a single plant family (Simon et al., 2015). For instance, the green peach aphid *Myzus persicae* attacks more than 400 plant species, whereas the cabbage aphid *Brevicoryne brassicae* and the pea aphid *Acyrthosiphon pisum* are specialized to *Brassicaceae* and *Fabaceae* families, respectively (Le Guigo et al., 2012; Silva et al., 2012). Such host specialization reflects their intimate relationship with host plants during their life cycle, which involves adaptations to cope with the plant phenology, the sap nutrient composition and chemical/physical defences. However, the aphid performance also depends on the interaction between its genotype or biotype and of the host-plant genotype or ecotype (Ferrari et al., 2008; Zytynska & Preziosi, 2013; Kanvil et al., 2014; Loxdale & Balog, 2018). During the last decade, the plant–microbe, arthropod–microbe and plant–arthropod interactions were shown to play an essential role in the selection, adaptation and evolution of plant–arthropod interactions, giving rise to the three-way interactions concept (Tétard-Jones et al., 2012; Biere & Bennett, 2013). While the effects of plant and aphid genotypes have been widely studied, the role of plant - rhizobia symbiosis has received less attention. Tétard-Jones et al., 2007 showed that interactions between barley genotypes and aphid clones are affected by the presence or absence of rhizospheric bacteria and evidenced the importance of the [aphid]x[barley]x[rhizobacteria] genotypes on aphid fitness in the multitrophic interactions. Recently, it was also suggested that plant growth-promoting rhizobia generally have negative effects on herbivore performance or abundance, most likely through Induced Systemic Resistance (ISR) in host plants (Pineda et al., 2010; Grunseich et al., 2020). However, the presence of rhizobium or arbuscular mycorrhiza (AM) fungi may have positive effects on sap-sucking insects likely due to an increase in phloem nitrogen or phosphorus levels (Dabré et al., 2022), and could also negatively affect herbivory natural enemies in the field, either by changing plant attractiveness or by decreasing the nutritive quality of their host (Pangesti et al., 2015; Peterson et al., 2016).

*Medicago truncatula* (barrel medic) is a model plant to explore leguminous plant biotic interactions. Multiple studies have shown that *M. truncatula* genotype plays a critical role in determining the interaction outcome with aphids that range from a fully compatible to a non-compatible plant-aphid interaction (Ferrari et al., 2008; Kanvil et al., 2014; Stewart et al., 2016). For instances, the *M. truncatula* Jester ecotype provides resistance against the blue-green aphid *Acyrthosiphon kondoi* and the pea aphid *A. pisum* Harris (Guo et al., 2008; Kamphuis et al., 2019), unlike the closely related ecotype A17, which gives no significant resistance to aphids (Klingler et al., 2005, 2007; Stewart et al., 2009). The variation in virulence among aphid genotypes on these different *M. truncatula* ecotypes led to the description of a molecular mechanism involving a ‘gene-for-gene’ recognition between resistance (R) genes in plants and associated avirulence (Avr) genes in aphids (Stewart et al., 2009; Kanvil et al., 2014). Similarly, variation in the induction of phytohormone-dependent plant response to aphid feeding depends on the plant and aphid genotypes. The blue green aphid induces the expression of genes associated with the salicylic acid (SA) pathway in both *M. truncatula* resistant and susceptible lines early after its attack, although with different induction kinetics (Gao et al., 2008). In contrast, genes associated with the jasmonic acid (JA) pathway are exclusively or predominantly induced in the resistant line (Gao et al., 2007, 2008). Moreover, *M. truncatula* can form an intimate nitrogen-fixing symbioses (NFS) with the bacteria *Sinorhizobium meliloti*, which not only affects plant growth but also modulates defence responses against herbivores (Dean et al., 2014; Franco et al., 2017). Pandharikar et al., 2020, demonstrated that NFS changes *M. truncatula* response to the pea aphid (*Acyrthosiphon pisum*) by affecting aphid growth and increasing the plant defence responses (Pandharikar et al., 2020). However, the study was conducted using a single plant and single aphid genotype. Yet, most of the previous works have examined only single plant–aphid genotype pairs, leaving the combined role of plant and aphid diversity under different symbiotic contexts largely unexplored.

Since both aphid and plant genotypes may contribute to the dynamics of these interactions, the present study aimed at unravelling genotype-dependent effects in this tripartite system based on three *A. pisum* genotypes and two *M. truncatula* ecotypes Jemalong A17 and R108. It has been demonstrated that the two *M. truncatula* ecotypes are different in their genomic structure, developmental traits, and physiological response to biotic and abiotic stresses (Salzer et al., 2004; Gaige et al., 2012; Li et al., 2014; Wang et al., 2014). We thus have quantified aphid fitness, plant biomass, and defence gene expression (SA and JA markers) in a time dependent manner to test how genotype-specific interactions and rhizobia inoculation jointly shape plant–aphid dynamics. We tested the hypothesized that (i) aphid performance would vary depending on plant genotype, (ii) rhizobia symbiosis would modulate defence expression and aphid success in a plant genotype-dependent manner, and (iii) outcomes would differ among aphid genotypes due to variation in virulence. The results demonstrate the critical role of genotypic diversity in influencing the ecological fate of multitrophic interactions and inform the complex relationship between plant nutritional status, symbiotic associations, and aphid pressure.

## 2. MATERIALS AND METHODS

### 2.1 Plant material and growth conditions

Two ecotypes of *M. truncatula*, Jemalong A17 and R108, were used. The two were chosen because they showed different resistance levels to pea aphids: A17 carries the *RAP1* gene, which provides race-specific resistance to the pea aphid (Kanvil et al., 2014). *RAP1* is ineffective against LL01 (alfalfa biotype) but effective against the pea and clover biotypes, while R108 has no resistance reported to date. Before sowing, *Medicago* seeds were treated (scarification, acid, etc.) as described (Barker et al., 2006; Nelson et al., 2015). The treated seeds were placed on 0.4% agar plates in the dark at 4°C for 2 days and then at 20°C for another 2 days. After germination, six plantlets were transplanted into round pots (7.5 x 7.5 cm) filled with a vermiculite and sand mixture (2:1). All pots were transferred to a growth chamber set at 23°C (16h light) and 20°C (8h dark), with a relative humidity of 60 ± 5%, and watered with nitrogen-free medium (Oger et al., 2012). One week after transplanting, *M. truncatula* A17 and R108 pots were divided into two groups: one group received inoculation with the *Sinorhizobium (Ensifer) meliloti* 2011 strain (named thereafter NFS plants) (del Giudice et al., 2011), and the second group was supplemented once with 10ml of 5 mM potassium nitrate (KNO_3_) in water (named thereafter NI plants) (Moreau et al., 2008). For inoculation, the *S. meliloti* 2011 strain was cultured on Luria-Bertani medium supplemented with 2.5 mM CaCl_2_ and MgSO4 (LBMC) and 200 μg/mL streptomycin for 3 days at 30°C. Subsequently, bacterial cells were grown in LBMC liquid medium for 24h, pelleted at 5000g, washed twice with sterile distilled water, and resuspended in sterile distilled water to a final optical density of 0.05 (OD 600) (Oger et al., 2012). Each NFS plant received a 10 mL supplementation of the *S. meliloti* suspension.

### 2.2 Aphids rearing and mesocosm preparation for aphid infestation

Three pea aphid (*A. pisum*) lines were used (Table S1): the original YR2 and T3-8V1 (abbreviated T38 thereafter) clones naturally harbour distinct strains of *Regiella insecticola*, and the removal of these symbionts through ampicillin treatment resulted in the creation of the YR2-amp and T3-8-V1-amp (named T38 hereafter) lines used here (Simon et al., 2011; Schmitz et al., 2012). LL01, in contrast, is naturally free of facultative symbionts. The YR2 and T38 lines are classified as "clover" biotypes, while LL01 is categorized as an "alfalfa" biotype. All aphid lines have been maintained stably for over 15 years on fava bean (*Vicia faba)*, at a temperature of 20°C under a 16:8h light/dark cycle (Fig S11). The absence of facultative symbionts was confirmed through PCR analysis (Luo et al., 2020), throughout the duration of the experiment. One week after inoculation with *S. meliloti* (allowing time to form nodules) or nitrate supplementation, NI and NFS pots (each containing six plants) were randomly assigned to different experimental groups and half of the pots were subjected to infestation with 10 first instar (L1) aphid nymphs (see Pandharikar et al., 2020). To synchronize aphid age, 20 wingless adult females were placed on separate fava bean plants, allowing them to reproduce for 24 hours. Then, 10 nymphs (L1) from each aphid line were transferred. All plant pots were individually isolated in ventilated plastic boxes and kept in the same room at 20°C under a 16:8h light/dark photoperiod with a 70% relative humidity.

### 2.3 Plant and aphid biomass

To assess the impact of aphid infestation on plant biomass (Fig S11), the dry weight of plant shoots was measured at 4, 8, and 12-days post-infestation (dpi). Plant shoots were dried in an 80°C oven for 3 days and subsequently weighed (balance accuracy ± 0.1 mg; PA214, Ohaus Corp, Parsippany, NJ, USA). The shoot dry weight of the six individual plants from three pots (total 18 plants) per condition was measured across three separate experiments. The development of the various pea aphid lines was assessed at 1, 4, 6, 8, and 12 days after aphid infestation (dpi). Aphid survival and their mean weight (total survived aphid weight/survived aphid number) were measured as fitness parameters. Six independent replicates were done for each time point, with distinct sets of plants and aphids. For each condition and time point, non-aphid infested NFS and NI plants were used as controls.

### 2.4 Gene expression analysis by quantitative RT-qPCR

Aphids were removed from the plants using paintbrush, and the control plants for each condition were brushed in the same way as aphid-infested plants. Six plant shoots per condition and time point were collected immediately after removal of the aphids, pooled and frozen in liquid nitrogen and store at −80°C when necessary. Three biological replicates were prepared for each condition and time point. For RNA extraction, the frozen shoots were grinded directly in liquid nitrogen using a mortar to obtain a fine powder. Total RNAs were isolated using RNAzol® RT (Sigma), quantified (NanoDrop 2000 spectrophotometer), and analyzed on a 1.5% agarose gel electrophoresis and a Bio-analyzer chips (Agilent) to assess purity. DNA digestion (RQ1 RNAse-free DNAse) and reverse transcription (GoScript™ Reverse Transcription) were performed as described by the manufacturer (Promega). The quantitative PCR was performed using the qPCR Master Mix plus CXR (qPCR kit; Promega). Each reaction was carried out with 5 μl of a 40-fold dilution of the cDNA template and each set of specific primers (Table S2). SA defence-related genes included *Medtr2g435490*, annotated as a *pathogenesis-related protein 1* (*PR.1*) (XP_013463163.1); *Medtr1g080800*, annotated as *chitinase (PR.4)*; and *Medtr1g062630*, annotated as *thaumatin-like protein* (*PR.5*) (XM_013612651.2). JA defence-related genes included *Allene Oxide Synthase 1* (*AOS.1*) (XM_013610584.2) and *Medtr4g032865*, a *proteinase inhibitor* (PSI-1.2) (XP_013455238.1), hereafter referred to as *PI*. Real-time qPCR was 95°C for 3 min followed by 40 cycles at 95°C for 5 sec and 60°C for 30 sec (AriaMx Real-time PCR machine, Agilent). The primers efficiency was evaluated on the slope of a standard curve generated using a serial dilution of the samples. Cycle threshold values (Ct) were normalized to the average Ct of two housekeeping genes (del Giudice et al., 2011): *Medtr2g436620*, which encodes the translationally-controlled tumor-like protein *MtC27* and is commonly used as a constitutive control in roots and nodules of *M. truncatula*, and *Medtr4g109650.1* (also known as a38), which encodes an uncharacterized protein. The original Ct values were obtained from the machine software (Ariamix software; Agilent) and further used in the RqPCRBASE package to normalize the expression level (Hilliou F & Tran T, 2013) using RStudio Version 1.1.453 (https://www.rstudio.com).

### 2.5 Replication statement

For clarity regarding the structure and replication of my dataset, a replication statement has been included with the manuscript.

### 2.6 Statistical analyses

For each of the nine dependent variables, the expression of the six genes, the plant dry weight, the aphid weight and survival, we used the following procedure to assess and test the interactive effect of genotypes and inoculation on their expression. We adjusted two mixed models with as fixed effect, the aphid genotype, the plant genotype, the plant inoculation, as well as the two- and three-way interactions between these variables. We also included as fixed effect the time variable (day of the experiment) as well as its two-way interactions with the aphid genotype, the plant genotype, the plant inoculation (the statistical power did not allow to assess higher order of the interactions here). We set the experimental batch and sub-batch as nested random effects. The two mixed models differ only for the time variable, which is a continuous variable in one case and a categorical variable in the other. The categorical one is better suited in case of non-linear relation with the dependent variable, but the number of parameters involved is also much higher. The best of these two models according to the Akaike Information Criterion (AIC) was used for further analysis. They were adjusted with the lme4 R package (Bates et al., 2015), and we used the DHARMa package (Hartig, 2020) to assess concordance between data and model assumptions, including checking for overdispersion in the GLMM model. The percentage of variation explained by the retained model was assessed by the marginal pseudo-R² (MuMIn package) (Bartoń, 2015; 2022) based on the likelihood-ratio test, with the null model containing the random effects: *marginal* pseudo-R² corresponds to percentage of variance explained by fixed effects only. When some significant interactions were found between the aphid genotype, plant genotype and plant inoculation, we interpreted them with post-hoc comparisons obtained by the multcomp package (Hothorn T et al., 2008). For each fixed effect, we estimated effect sizes using partial-η² (Grissom & Kim, 2005), computed using explained sum of squares (ESS) and residual sum of squares (RSS) estimated using type 2 decomposition of variance (partial-η²=ESS/(ESS+RSS). As stated by Cohen 1988 (Cohen J, 1988), it is a proportion, not of the total variance, but of the total from which there has been excluded the variance due to the other factor(s) and interactions. In order to ease their interpretation, we categorized these effect sizes using the thresholds proposed by Cohen 1988 and Richardson 2011(Effect size: 0.0099 < Small < 0.0588 < Medium < 0.1379 < Large).

## 3 RESULTS

We used a three-way interaction model to explain how plant biomass, aphid fitness, and plant defence gene expression are influenced either by individual factors or by interactions among plant genotype, aphid genotype, and plant rhizobium inoculation during the time frame of the study.

### 3.1 Aphid infestation negatively impact plant biomass independent of genotype or inocu-lation

At the start of the experiment, the shoot dry mass of the NI and NFS plants was similar (Fig S1). Over the 12 days, the dry weight of the control plants was affected by the plant genotype (partial η² = 0.18, *P* < 0.001) and the nitrogen source (partial η² = 0.446, *P* < 0.001) (Fig S2, Table 1).

**Table 1.** Linear mixed-effects models assessing the effects of aphid genotype, plant genotype, inoculation with NFS or NI, and time (categorical) on plant dry weight. For each factor and interaction, η² values (effect size estimates), their interpretation (small, medium, large), and statistical significance are reported. Significance levels are indicated as follows: ***p < 0.001; **p < 0.01; *p < 0.05; p < 0.1; ns = not significant. Effect size interpretations follow standard thresholds (η²: small ≥ 0.01, medium ≥ 0.06, large ≥ 0.14). Significance levels are indicated as follows: ***p < 0.001; **p < 0.01; *p < 0.05; p < 0.1 ·p < 0.1 (marginal significance).

| Factor | $\eta^2$ | Effect Size | Significance |
| --- | --- | --- | --- |
| Aphid G. | 0.5 | Large | < 0.001 *** |
| Plant G. | 0.18 | Large | < 0.001 *** |
| Inoculation | 0.45 | Large | < 0.001 *** |
| Time | 0.5 | Large | < 0.001 *** |
| Aphid G. × Plant G. | 0.02 | Small | < 0.001 *** |
| Aphid G. × Inoculation | 0.068 | Medium | < 0.001 *** |
| Plant G. × Inoculation | 0.026 | Small | < 0.001 *** |
| Aphid G. × Time | 0.18 | Large | < 0.001 *** |
| Plant G. × Time | 0.059 | Medium | < 0.001 *** |
| Inoculation × Time | 0.13 | Medium | < 0.001 *** |
| Aphid G. × Plant G. × Inoculation | 0.0041 | Not specified | 0.33 (ns) |

As expected, aphid infestation had a strong impact on plant dry weight (partial η² = 0.5, *P* < 0.001), and this was independent of the plant nutritional status (NI or NFS) (Figure S2; S7-10). While significant interactions were found between the plant genotype and the aphid genotype (partial-η^2^ =0.02, p < 0.001), aphid genotype x plant inoculation (partial-η^2^ =0.068, p < 0.001) and plant genotype x plant inoculation (partial-η^2^ =0.026, p < 0.001), the three-way interaction plant inoculation x aphid genotype x plant genotype was not significant (p = 0.33) (Table 1). Together, these results show that aphid infestation consistently reduces plant biomass regardless of nitrogen source, while the scale of this reduction was further shaped by specific plant–aphid–*S. meliloti* inoculation combinations.

### 3.2 Plant, aphid genotypes and plant inoculation modulate aphid weight

Aphids successfully survived and developed on both A17 and R108 *M. truncatula* genotypes regardless of the nitrogen nutritional source. However, some aphid genotypes grew better on A17 genotype and others on R108 genotype, as shown by a large effect (partial η² = 0.32, P < 0.001) on aphid weight, and by the large interaction between aphid genotype x plant genotype (partial η² = 0.42, P < 0.001) (Fig 1, Fig S6, Table 2).

**Figure 1.**
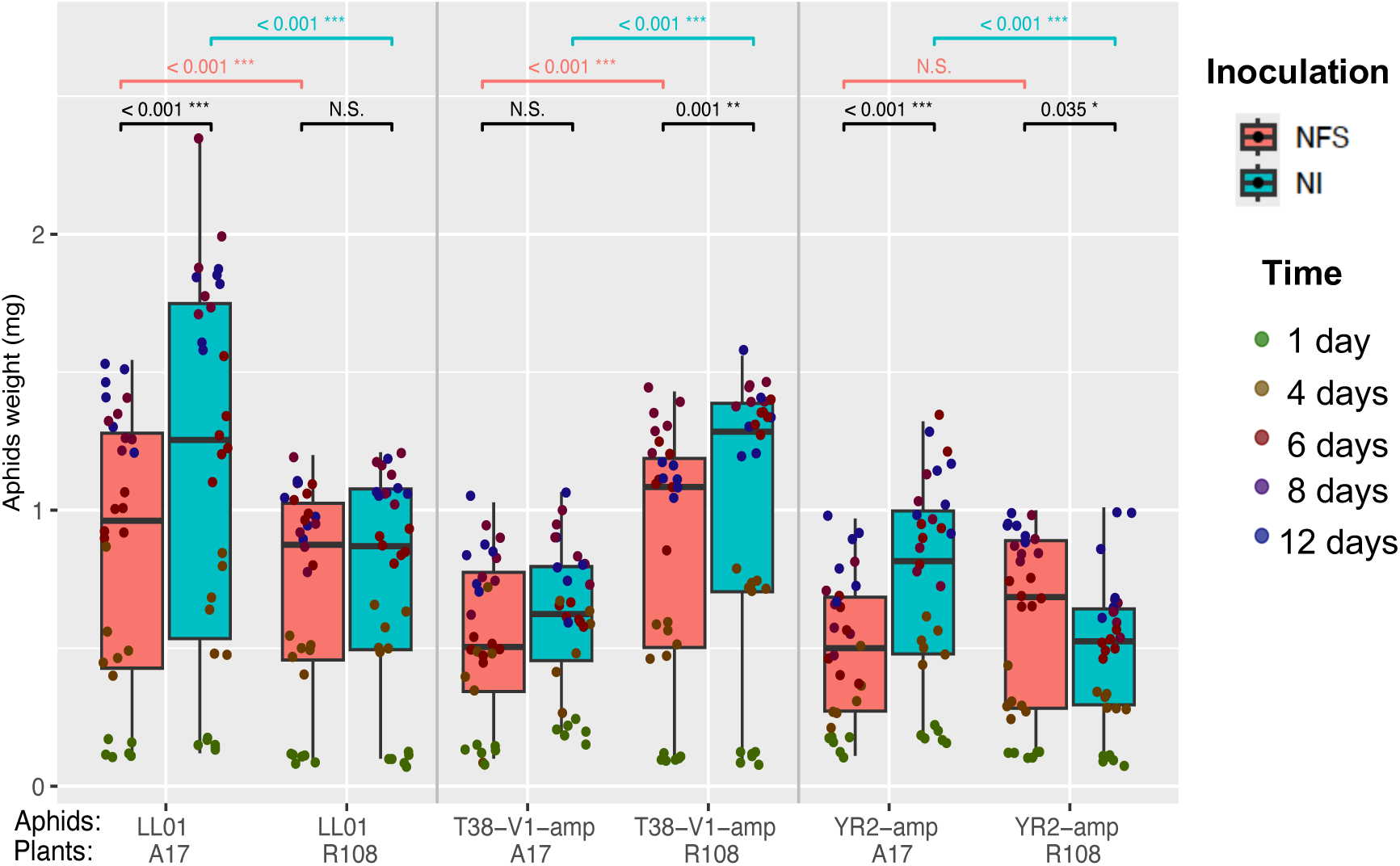
Aphid weight over time on different *M. truncatula* genotypes and inoculation treatment. Box plots showing the mean weight of the surviving aphid (*A. pisum* clones LL01, YR2-amp, and T38-V1-amp) on two *Medicago truncatula* genotypes (A17 and R108) under two nitrogen treatment: mineral nitrate supplementation (NI in blue) and rhizobia inoculation (*Sinorhizobium meliloti*; NFS in pink). Aphid weight was measured at five independent time points (1, 4-, 6-, 8-, and 12-days post-infestation; *n* = 6 per time point). The X-axis legend indicates aphid–plant genotype combinations. Jittered points represent individual biological repeat for mean weight of the living aphids, with point colours indicating the sampling time. Mixed model predictions for the interaction of plant inoculation, aphid and plant genotype on aphid weight (Marginal R² = 0.90)

**Table 2.** Linear mixed-effects models assessing the effects of aphid genotype, plant genotype, inoculation with NFS or NI, and time (categorical) on aphid weight (fitness). For each factor and interaction, η² values (effect size estimates), their interpretation (small, medium, large), and statistical significance are reported. Significance levels are indicated as follows: ***p < 0.001; **p < 0.01; *p < 0.05;. p < 0.1; ns = not significant. Effect size interpretations follow standard thresholds (η²: small ≥ 0.01, medium ≥ 0.06, large ≥ 0.14). Significance levels are indicated as follows: ***p < 0.001; **p < 0.01; *p < 0.05; p < 0.1 ·p < 0.1 (marginal significance).

| Factor | $\eta^2$ | Effect Size | Significance |
| --- | --- | --- | --- |
| Aphid G. | 0.32 | Large | < 0.001 *** |
| Plant G. | 0.055 | Small | < 0.001 *** |
| Inoculation | 0.13 | Medium | < 0.001 *** |
| Time | 0.79 | Large | < 0.001 *** |
| Aphid G. × Plant G. | 0.42 | Large | < 0.001 *** |
| Aphid G. × Inoculation | 0.017 | Small | 0.059 . |
| Plant G. × Inoculation | 0.087 | Medium | < 0.001 *** |
| Aphid G. × Time | 0.26 | Large | < 0.001 *** |
| Plant G. × Time | 0.045 | Small | 0.005 ** |
| Inoculation × Time | 0.028 | Small | 0.053 . |
| Aphid G. × Plant G. × Inoc. | 0.095 | Medium | < 0.001 *** |

For instance, LL01 aphid genotype gained significantly higher weight on A17 than R108 (Fig 1), whereas T38 genotype performed significantly better on R108 than A17 across both nitrogen regimes (Fig 1). YR2 grew better on A17 than R108 under NI feeding (Fig 1), but this difference disappeared under NFS conditions. Plant rhizobium inoculation (NFS) had a medium effect (partial η² = 0.13, P < 0.001) on aphid weight, and interacted close to significance with aphid genotype (partial η² = 0.017, P = 0.059). For example, NFS reduced significant aphid weight in several host-aphid combinations (LL01/A17, T38-/R108, YR2/A17) (Fig 1), but had no significant effect on LL01/R108 or T38/A17 (Fig 1), and even significantly enhanced growth on YR2/R108 (Fig 1, Fig S6). The three-way interaction between aphid genotype, plant genotype, and plant inoculation had a medium effect (partial η² = 0.095, P < 0.001) (Table 2, Fig 1). Overall, these results showed that aphid fitness was strongly influenced by plant–aphid genotype interactions and further modulated by rhizobia symbiosis.

### 3.3 Plant and aphid genotypes and plant inoculation modulate defence gene expression

Expression levels of *PR.1*, *PR.4* and *PR.5* were influenced by both plant and aphid genotype, in a time dependent manner (Table S3). Compared to control (non-infested) plants, over the time, aphid infested plants showed significantly higher gene expression levels of SA-mediated marker genes (Fig 2; S3-S4). Aphid genotype had large effect on *PR.1* (Fig 2, Table 3) (partial η² = 0.19, *P* < 0.001) and *PR.5* expression (partial η² = 0.23, *P* < 0.001), while the effect on *PR4* was medium (partial η² = 0.065, *P* = 0.015) (Fig S3-S4; Table S3). Plant genotype had a medium effect on *PR.4* (partial η² = 0.11, *P* < 0.001) but was not significant for *PR.1* (η² = 0.0038, *P* = 0.53) or *PR.5* (η² = 7e-4, *P* = 0.79) (Table S3). Plant rhizobia inoculation (NFS) resulted in a medium effect on *PR.5* expression (partial η² = 0.12, *P* < 0.001), whereas its effect on *PR.1* and *PR.4* was non-significant (*PR.1* η² = 0.0061, *P* = 0.4; *PR.4* η² = 0.012, *P* = 0.23) (Table 3, Fig S4, Fig S5). The three-way interaction between aphid genotype, plant genotype, and plant inoculation significantly influenced the expression of *PR.1* (*η²* = 0.08, *P* = 0.008) but not *PR.4* and *PR.5* (Table S3, Fig S4, Fig S5).

**Figure 2.**
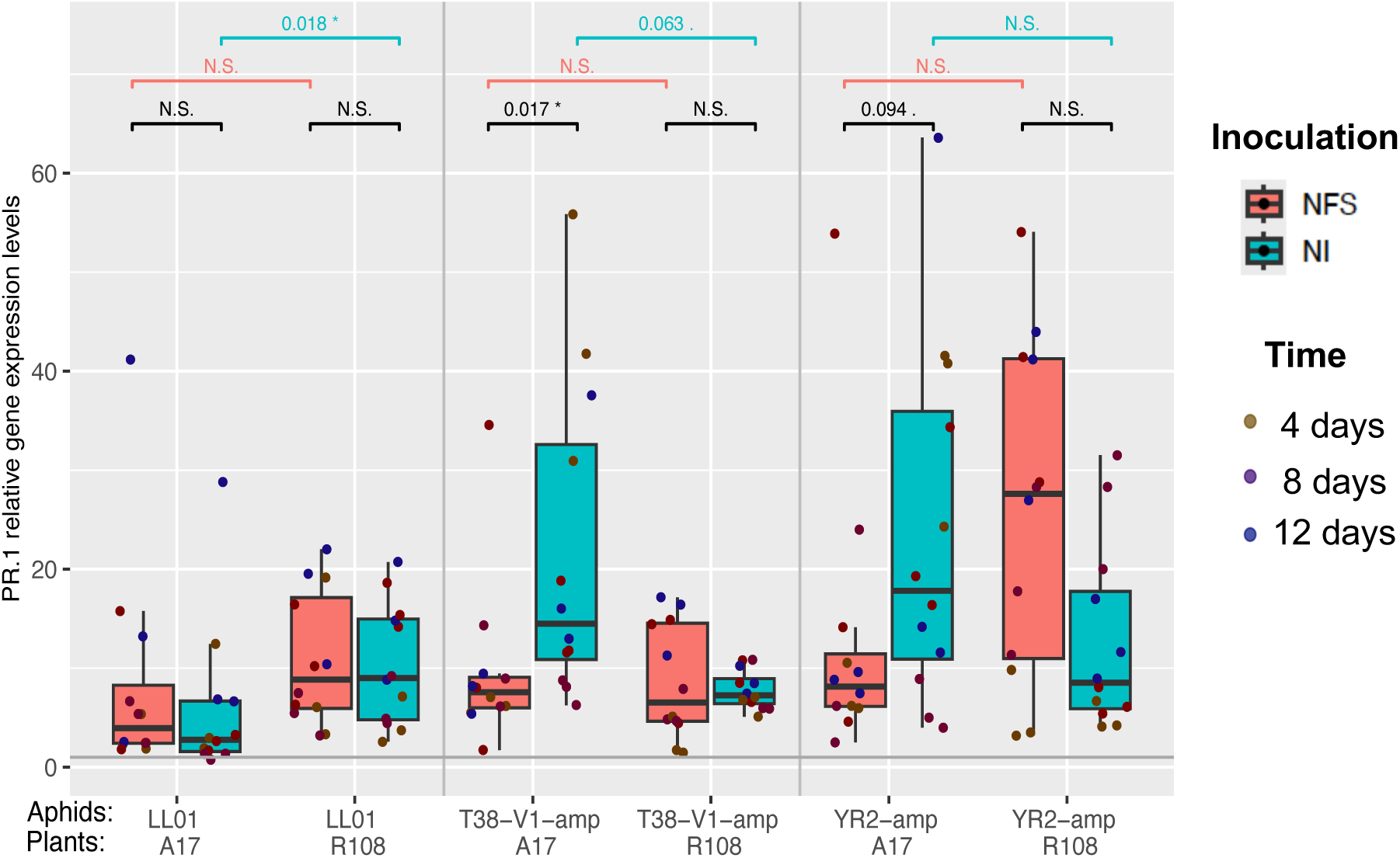
*PR.1* gene expression in *Medicago truncatula* shoots following aphid infestation and inoculation treatment. Expression levels of the salicylic acid (SA) pathway marker gene *PR.1* in shoots of two *Medicago truncatula* genotypes (A17 and R108) infested with three aphid genotypes (LL01, YR2-amp, T38-V1-amp), under two nitrogen treatments: nitrate-fed (NI; blue) and rhizobia-inoculated (NFS; pink) (n=3; each replicate a pool of 6 plants). Expression values are rescaled relative to non-infested control plants (grey horizontal line fitted at 1), which represent basal expression levels. Box plots show median and interquartile range (IQR); dots coloured represent individual biological replicate for *PR.1* gene expression levels observations at discrete time points post-infestation. Mixed model predictions for the interaction of plant inoculation, aphid and plant genotype on *PR.1* gene expression (Marginal R² = 0.57)

**Table 3.**
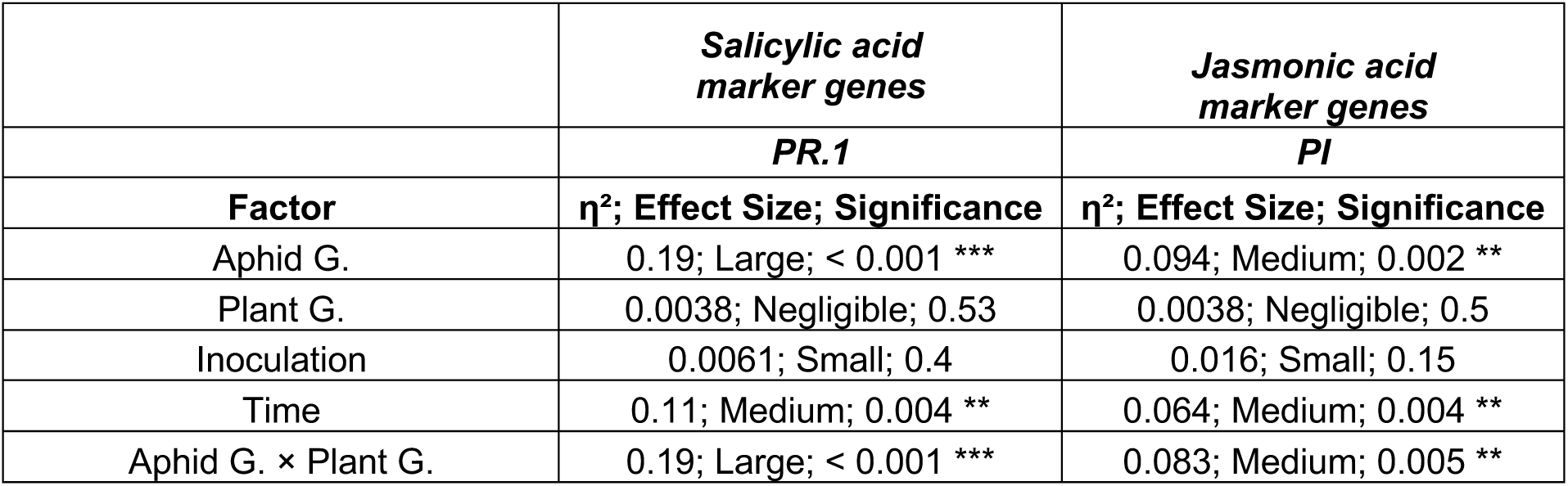
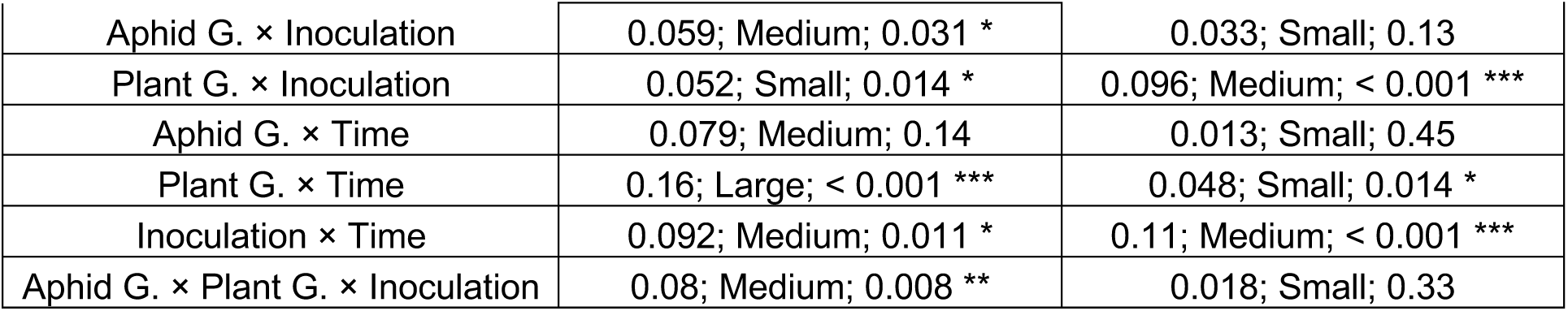
Linear mixed-effects models assessing the effects of aphid genotype, plant genotype, inoculation with NFS or NI, and time (categorical) on salicylic acid marker genes (*PR.1)* and Jasmonic acid marker genes (*PI*). For each factor and interaction, η² values (effect size estimates), their interpretation (small, medium, large), and statistical significance are reported. Significance levels are indicated as follows: ***p < 0.001; **p < 0.01; *p < 0.05; p < 0.1; ns = not significant. Effect size interpretations follow standard thresholds (η²: small ≥ 0.01, medium ≥ 0.06, large ≥ 0.14). Significance levels are indicated as follows: ***p < 0.001; **p < 0.01; *p < 0.05; p < 0.1 ·p < 0.1 (marginal significance).

Expression of Jasmonic acid (JA) pathway marker genes, *AOS.1* (JA biosynthesis) and protease inhibitor (*PI*) (JA responsive gene), was regulated by genotype, inoculation and time factors (Table S3, Fig 3, Fig S4, Fig S5). For *PI*, aphid genotype had a medium effect (η² = 0.094, p = 0.002), plant inoculation (η² = 0.016, p = 0.15) and plant genotype alone did not significantly influence *PI* expression (η² = 0.0038, p = 0.5) (Table 3, Fig 3). However, interactions between plant genotype and inoculation (η² = 0.096, p < 0.001), and between aphid genotype and plant genotype (η² = 0.083, p = 0.005), had medium effect (Table 3). Time also significantly affected *PI* expression (η² = 0.064, p = 0.004), and its interaction with plant inoculation was particularly strong (η² = 0.11, p < 0.001) (Table 3). For *AOS1*, only inoculation produced a medium effect (η² = 0.085, p = 0.001) (Table S3, Fig S5) and no significant effect of time was detected (η² = 0.014, p = 0.65). In summary, defence gene expression in both SA- and JA-pathways was differentially regulated by aphid genotype, plant genotype, and rhizobia inoculation, with several significant two and three-way interactions detected.

**Figure 3.**
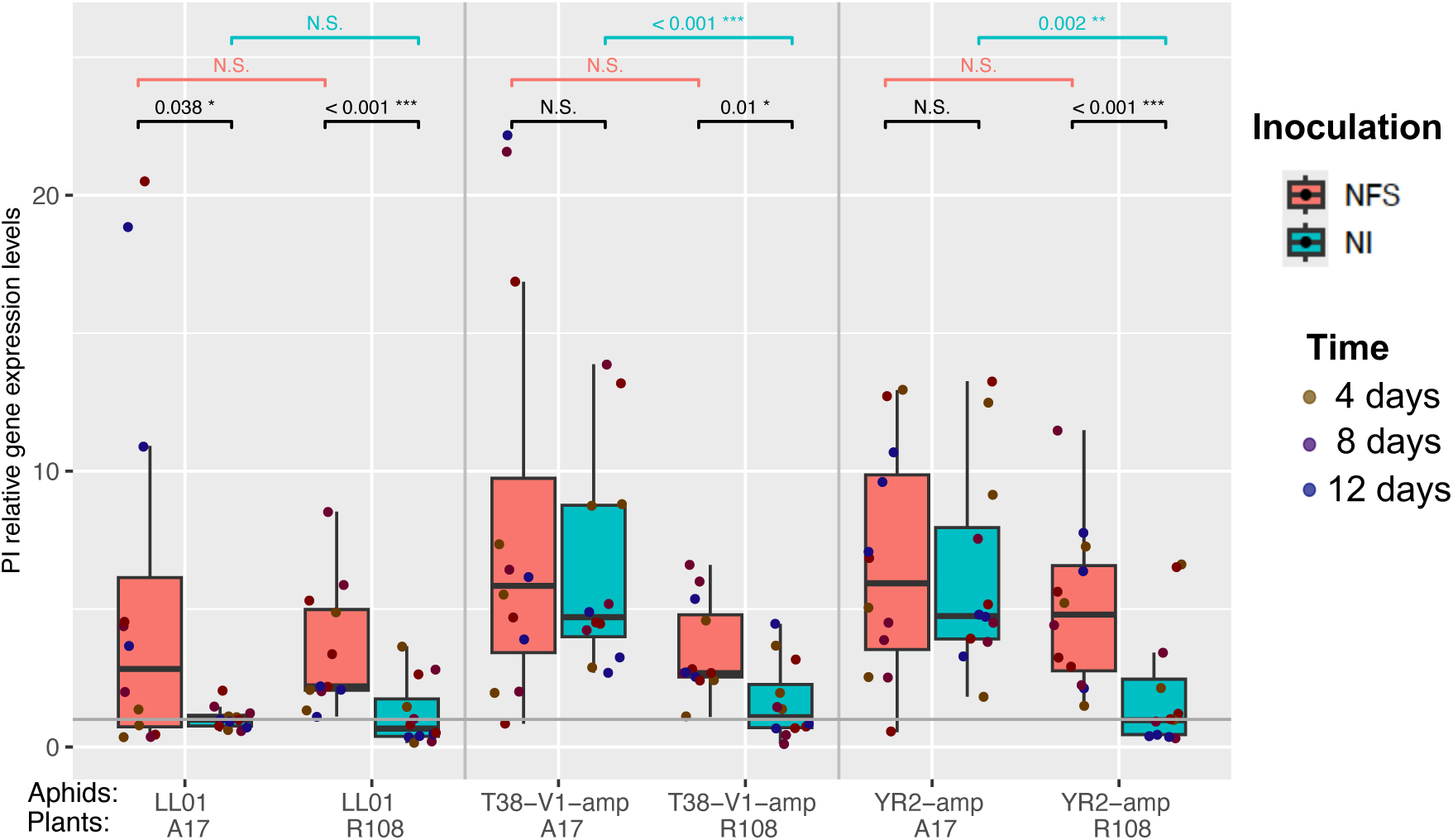
*PI* gene expression in *Medicago truncatula* shoots following aphid infestation and inoculation treatment. Expression levels of the Jasmonic acid (JA) pathway marker gene *PI* in shoots of two *Medicago truncatula* genotypes (A17 and R108) infested with three aphid genotypes (LL01, YR2-amp, T38-V1-amp), under two nitrogen treatments: nitrate-fed (NI; blue) and rhizobia-inoculated (NFS; pink) (n=3; each replicate a pool of 6 plants). Expression values are rescaled relative to non-infested control plants (grey horizontal line fitted at 1), which represent basal expression levels. Box plots show median and interquartile range (IQR); dots represent individual biological replicate for *PI* gene expression levels observations at discrete time points post-infestation. Mixed model predictions for the interaction of plant inoculation, aphid and plant genotype on *PI* gene expression (Marginal R² = 0.5).

## Discussion

The genetic interactions between aphids and legumes are classically viewed as determined by the genotypes of both organisms, producing either compatible or incompatible outcomes (Goh et al., 2013; 2016; Kanvil et al., 2014). Aphid virulence appears to be a complex trait involving multiple genes that influence factors such as saliva composition, which in turn modulates plant defence responses and host compatibility (Thorpe et al., 2024; Ollivier et al., 2025). Legume plants also form symbiotic relationships with nitrogen-fixing rhizobia, which can transiently activate early plant defence reactions that influence susceptibility to pathogens (Lohar et al., 2003; Grundy et al., 2023). However, few studies have examined how these plant-rhizobia symbiosis intersect with the genotype of both plant and aphid to shape defence and aphid fitness outcomes. To investigate how this interaction (plant genotype, aphid genotype, and rhizobia symbiosis) shapes outcomes, we studied two *M. truncatula* cultivars, A17, carrying the aphid resistance QTL RAP1 (Stewart et al., 2009), and R108, which lacks RAP1, and appears more susceptible with the three pea aphid lines representing different biotypes (LL01, an "alfalfa" biotype; YR2 and T38, "clover" biotypes). Our analysis focused on three key aspects: plant growth, aphid fitness, and plant defence responses during co-development of aphids and rhizobia nodules (see also Pandharikar et al., 2020). This comprehensive approach helps to unravel how plant-rhizobia symbiosis modulates both direct plant resistance mechanisms and indirect effects on aphid success, considering the genotypic interplay across all three partners.

### Genotypes and symbiotic status modify plant biomass after aphid infestation

Over the 12 days experimental time frame, the nitrate fed (NI) plants accumulate a significantly higher shoot dry biomass compared to those in rhizobia symbiosis (NFS) (Fig S2, Fig S7, Fig S10). This was expected since nitrate provides immediate and readily available nitrogen, whereas symbiotic nitrogen fixation requires significant carbon and energy investment to establish nodules to reach the maximum of nitrogen compounds availability (Goh et al. 2013; Pandharikar et al. 2020; Wan et al., 2025). A strong effect of the plant genotype x inoculation was also observed, may be due to the fact that A17 and R108 have different symbiotic effectiveness with different *Sinorhizobium* (Ensifer) inoculum (Kazmierczak et al., 2017; Burghardt et al., 2018). At 21 days post inoculation, A17 has been reported to form more effective nodules with *S. meliloti* 2011 than R108 (Luu et al., 2022). Similarly, our results show that NFS R108 plants had lower shoot biomass compared to NFS A17, which could be due to reduced nitrogen availability due to fewer or less efficient nodules. Moreover, Pandharikar et al., 2020, along with other studies, have shown that aboveground aphid attacks can affect legume–rhizobia symbiosis by reducing nodule formation (e.g., decreases in nodule size, number, and weight, as well as dry biomass) and by downregulating the expression of genes involved in various steps of nodule development, ultimately leading to reduced nitrogen fixation (Pandharikar et al., 2020; Basu et al., 2021). As expected, infestation by aphid reduced shoot plant dry weight in both NI and NFS treated (Fig S2, Fig S7, Fig S10) since aphid feeding acts as a carbon sink hijacking resources from the plant (Girousse et al., 2005). Similar biomass loss independent of rhizobia status have been reported with *Aphis glycines* on soybean (Glycine max L.) (Brunner et al., 2015) and is consistent with Pandharikar et al. 2020. The extent of biomass reduction was not uniform but depended on specific plant–aphid genotype combinations. Using 8 aphid lines including LL01, Kanvil et al., 2014 showed that aphid lines differ in their ability to induce chlorosis and necrosis across A17 and R108. Chlorosis in R108 was common across multiple aphid lines, while necrosis was observed in A17 but not R108 (Kanvil et al., 2014; Stewart et al., 2016). This chlorosis associated to hypersensitive response was not quantified in our study, but chlorosis may affect photosynthesis and then plant growth, reducing the plant biomass of R108 in addition to the direct effect of aphid feeding. The extent of biomass loss depended on specific plant-aphid genotype combinations, supporting that genotype of plant and aphid critically influences aphid-driven plant growth inhibition.

### Context dependent modulation of aphid fitness by plant genotype and rhizobia

All aphid lines survived equally on both *M. truncatula* A17 and R108 genotypes but showed significant differences in aphid weight (a proxy for fitness) (Fig 1) driven by the specific plant–aphid genotype and rhizobia pairing. LL01 displayed consistently higher weight on A17 across all conditions (NFS and NI) (Fig 1), consistent with prior reports that LL01 expresses broad virulence and is specifically virulent to overcome A17 defences (Kanvil et al., 2014). Conversely, when LL01 fed on R108, its fitness significantly dropped, indicating that R108 presents a stronger or more specific defense to LL01, restricting its compatibility range. T38 aphid genotype performed better on R108 than on A17, suggesting it might possess virulence traits or physiological adaptations that target the defence response present in R108, but are less effective on A17. The third genotype, YR2, had comparatively low fitness across both plant hosts, implying that it is less virulent or has a narrower host adaptation and thus, encounters robust resistance regardless of host. These results indicate how diversity of plant and aphid genotypes and gene expression allows differential exploitation of host resources, a key strategy for persistence in different environments (Stewart et al., 2016; Eyres et al., 2016). Plant rhizobia inoculation further modify aphid fitness, with NFS often reducing aphid weight (LL01 on A17; T38 on R108; YR2 on A17) (Fig 1), but in some combinations (LL01 on A17; T38 on R108) inoculation improves aphid performance (YR2 on R108). These results suggest that plant symbiotic status (NFS) can sometime counterbalance the nutritional advantage of NI and that the aphid growth rate is not linked directly to the plant growth rate. These results also underscore that rhizobia inoculation does not uniformly increase host nutritional value for aphids, but instead reshapes host quality in a genotype-dependent manner. This agrees with studies showing that rhizobia alter phloem sap amino acid composition, sugar ratios, and secondary metabolite levels, which in turn affect herbivore performance (Zytynska & Weisser, 2016; Gualtieri et al., 2021). For instance, enhanced nitrogen assimilation via rhizobia may increase essential amino acids in phloem sap, benefiting some aphid genotypes, while simultaneous induction of phenolics or terpenoids may reduce fitness in others. Recent work suggests that the net effect of rhizobia inoculation is conditional, producing either induced resistance or induced susceptibility depending on the host–aphid genetic pairing (Zhong, 2019; Dabré, 2022).

### Genotype-specific and inoculation-driven modulation of plant defence responses

The plant defence signalling in response to aphid infestation was strongly influenced by plant × aphid genotype (G×G) interactions, while rhizobia inoculation introduces an important layer of modulation. This dual influence shows how defence outcomes are not simply add-on, but emerge from a dynamic interplay between host genotypes and microbial partners. Previous studies have shown that aphid populations differ in their ability to induce or suppress plant defence signalling pathways based on G×G interactions with host plant (Giordanengo et al., 2010; Jaouannet et al., 2014). This aligns with our observation that aphid lines such as T38 and YR2 caused a significant downregulation of *PR.1* a marker of salicylic acid (SA)-mediated defence in Medicago genotype A17 (NI), but not in R108 (NI) (Fig 2). The contrasting reaction of R108 suggests that host genetic background constrains or enables aphid virulence factors to manipulate defence signalling. Such genotype × genotype (G×G) dynamics represent a molecular arms race, where host resistance traits and aphid effectors co-evolve to yield highly specific response patterns (Glazebrook 2005; Stout et al., 2006). Importantly, these G×G outcomes were not limited to SA pathways. We also observed variable regulation of the jasmonic acid-associated *PI* gene (Fig 3), which was upregulated in R108 under NFS conditions with all aphid genotype, but only triggered by the LL01 aphid genotype in A17 (NFS). This differential *PI* regulation suggests that nodulation capacity and host–rhizobia compatibility influence the disbilirection of defence modulation. This supports the idea that microbial partners can reprogram the defence status of plants, sometimes enhancing resistance, other times creating vulnerabilities exploitable by aphids (Glazebrook 2005; Stout et al., 2006; Benjamin et al., 2024). NFS plants showed genotype-specific shifts in the expression of both SA- and JA-related genes. This supports the hypothesis that symbiosis with nitrogen-fixing bacteria alters the plant immune status, either through direct modulation of hormone balances or via systemic resource reallocation (Zytynska & Preziosi 2013; Zytynska et al., 2014; Sanchez-Arcos et al., 2016). Recently, Mbaluto & Zytynska 2025 showed that Rhizobacteria inoculation suppress aphids on barley by priming plant defense and nutritional pathways. They enhance resistance plant resistance either through phenylpropanoid, glutathione, and phytohormone pathways, while also promoting tolerance via improved nutrition and growth, highlighting a dynamic, multi-pathway mechanism of microbial-induced plant protection (Mbaluto & Zytynska 2025).

Mechanistically, rhizobial colonization is known to induce cross-talk between nodulation pathways and phytohormone signalling, in particular SA, JA, and Ethylene pathways. Such cross-talk can dampen SA-dependent resistance to pathogens or herbivores while enhancing JA-dependent responses in some contexts (Sanchez-Arcos et al., 2016). In our study, the observed genotype-dependent defence shifts suggest that the inoculation effect is not uniform but instead depends on the plant host basal capacity to integrate both nodulation and defence signalling. Thus, rhizobia represent a conditional “third player” in the plant–aphid arms race, capable of swinging the defensive balance in favour of or against the plant, depending on the genetic pairing of both plant and aphid. Overall, our results reflect a broader principle of multitrophic interactions, where symbiotic microbes such as rhizobia can recalibrate plant immune response, altering outcomes of ongoing interactions with aphids.

## Conclusion

Our findings highlight that plant–aphid interactions are shaped not only by the genetic identity of the interacting species but also by the plant symbiotic context. Genotype-specific responses in both defence signalling and aphid performance underscore the ecological importance of variation within and between species. Importantly, rhizobia symbiosis emerged as a key modulator of plant defence and growth, reinforcing the idea that microbial mutualists must be integrated into ecological and evolutionary models of plant resistance. By showing that microbial associations can either enhance or constrain herbivore resistance depending on host genetic background, our findings open new avenues for exploiting beneficial symbioses to strengthen crop resilience. Targeted use of plant genotype–microbe combinations could provide a sustainable strategy to improve plant defence against insect pests while maintaining ecosystem functions.

## Acknowledgements

We are grateful to the ISA symbiosis team (IRL) members for their help and assistance in the experimental work. We also acknowledge Dr S. Tares and L. Arthaud for their help with aphid rearing and the MIB team members for constant help and support.

## Data accessibility

The complete raw dataset and corresponding analysis scripts can be accessed at following linkhttps://entrepot.recherche.data.gouv.fr/dataset.xhtml?persisten-tId=doi:10.57745/WGBWYO

## Authors contributions

G.P., M.P., P.F., J.-L.G.: experimental design and conceptualization; G.P.: data acquisition and original draft preparation; H.M.H: statistical analysis; J.-C.S.: supply of aphid biological resources; G.P., J.-C.S., J.-L.G., and P.F: writing-review and editing. M.P., P.F., J.-L.G.: supervision and funding acquisition.

## Supplementary Figures

**Figure S1:**
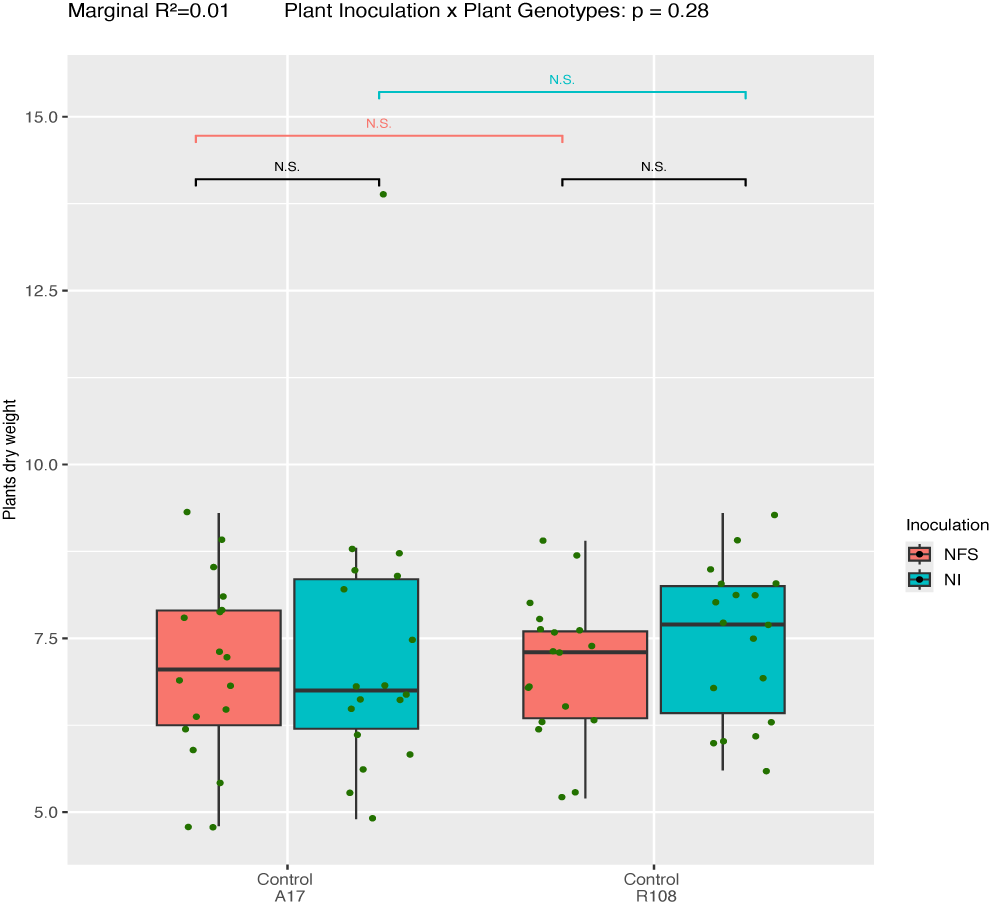
Comparison of the dry weight of the NFS and NI plants of both A17 and R108 *Medicago truncatula* genotype before aphid infestation. Box plots show shoot dry weight of *Medicago truncatula* (genotypes A17 and R108) obtained at 14 days after seeding and 7 days after Kno3 or Rhizobia treatments; either watering with a mineral nitrate solution (NI) or inoculation with *Sinorhizobium meliloti* (NFS). Boxes represent medians and interquartile ranges; Pink boxplots represent the NFS condition, while blue boxplots represent the NI condition. (n=3; each replicate a pool of 6 plants); N.S. = not significative.

**Figure S2:**
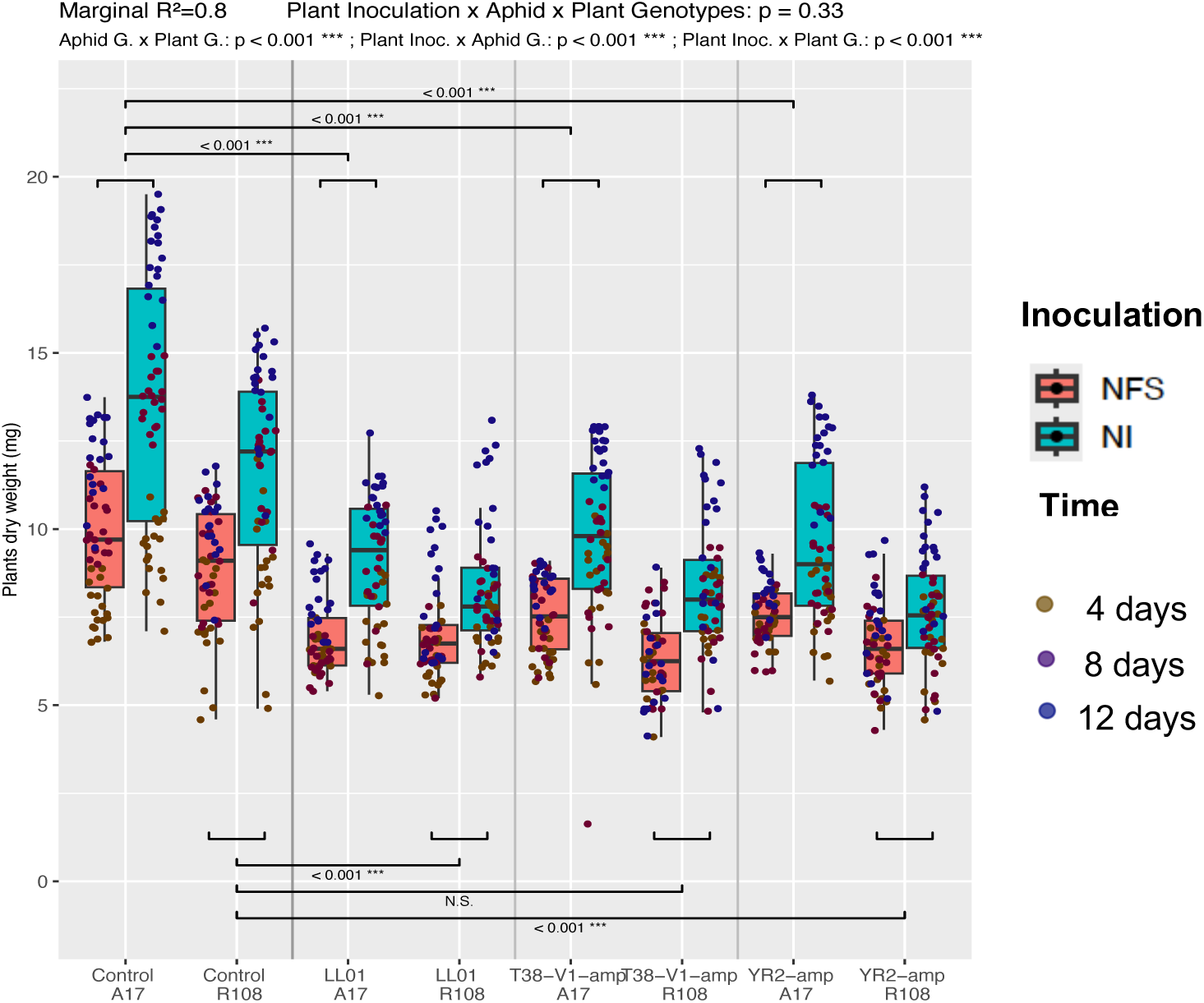
Effects of plant genotype, aphid genotype, and rhizobia inoculation on dry weight in *Medicago truncatula*. Box plots show shoot dry weight of *Medicago truncatula* (genotypes A17 and R108) exposed to three pea aphid clones (LL01, YR2-amp, T38-V1-amp) under two nitrogen treatments: mineral nitrate supplementation (NI) or inoculation with *Sinorhizobium meliloti* (NFS) (n=3; each replicate a pool of 6 plants). Boxes represent medians and interquartile ranges; Pink boxplots represent the NFS condition, blue boxplots represent the NI condition. The jitter dots represent each individual plant for dry weight measurements at discrete time points post-infestation (see time colours on the legend). Mixed model predictions for the interaction of plant inoculation, aphid and plant genotype on dry weight (Marginal R² = 0.8)

**Figure S3.**
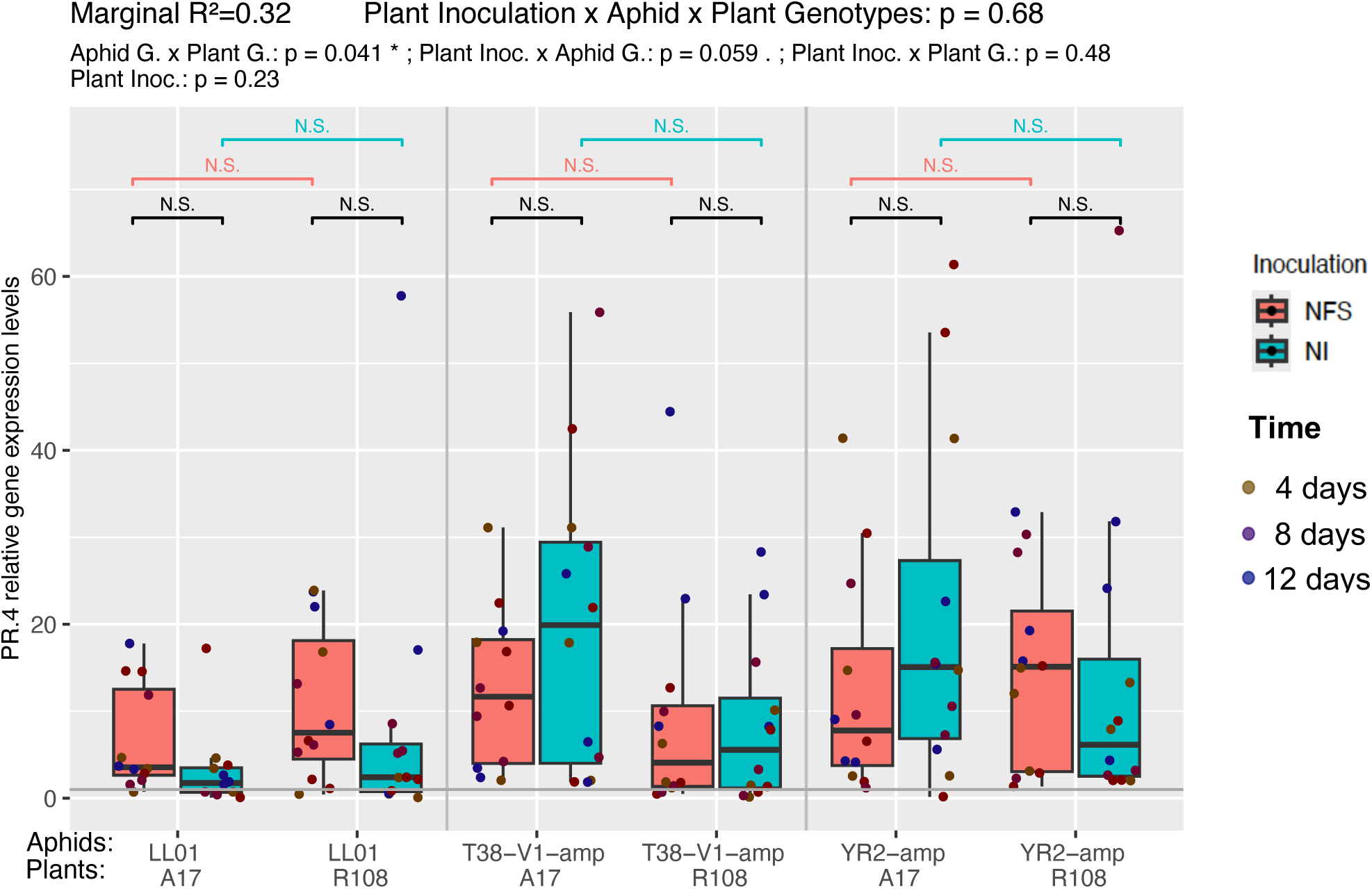
*PR.4* gene expression in *Medicago truncatula* shoots following aphid infestation and inoculation treatment. Expression levels of the salicylic acid (SA) pathway marker gene *PR.4* in shoots of two *Medicago truncatula* genotypes (A17 and R108) infested with three aphid genotypes (LL01, YR2-amp, T38-V1-amp), under two nitrogen treatments: nitrate-fed (NI; blue) and rhizobia-inoculated (NFS; pink) (n=3 (each replicate a pool of 6 plants)). Expression values are rescaled relative to non-infested control plants (grey horizontal line), which represent basal expression levels. Box plots show median and interquartile range (IQR); dots represent individual biological replicate for *PR.4* gene expression levels measurements at discrete time points post-infestation (see time colours on the legend). Mixed model predictions for the interaction of plant inoculation, aphid and plant genotype on *PR.4* gene expression (Marginal R² = 0.32).

**Figure S4.**
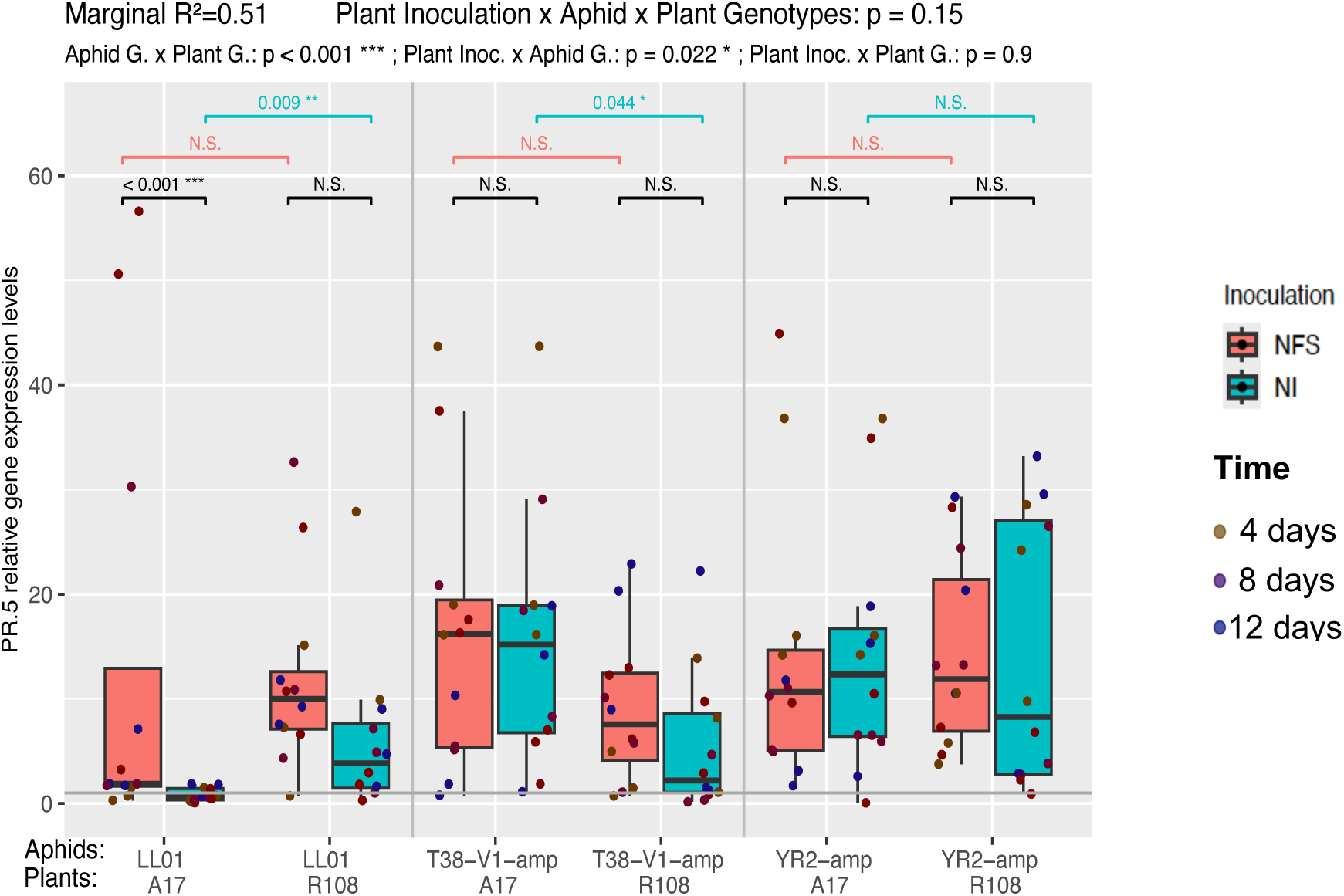
*PR.5* gene expression in *Medicago truncatula* shoots following aphid infestation and inoculation treatment. Expression levels of the salicylic acid (SA) pathway marker gene *PR.5* in shoots of two *Medicago truncatula* genotypes (A17 and R108) infested with three aphid genotypes (LL01, YR2-amp, T38-V1-amp), under two nitrogen treatment: nitrate-fed (NI; blue) and rhizobia-inoculated (NFS; pink) (n=3; each replicate a pool of 6 plants). Expression values are rescaled relative to non-infested control plants (grey horizontal line), which represent basal expression levels. Box plots show median and interquartile range (IQR); dots coloured represent individual biological replicate for *PR.5* gene expression levels observations at discrete time points post-infestation (see time colours on the legend). Mixed model predictions for the interaction of plant inoculation, aphid and plant genotype on *PR.5* gene expression (Marginal R² = 0.51)

**Figure S5.**
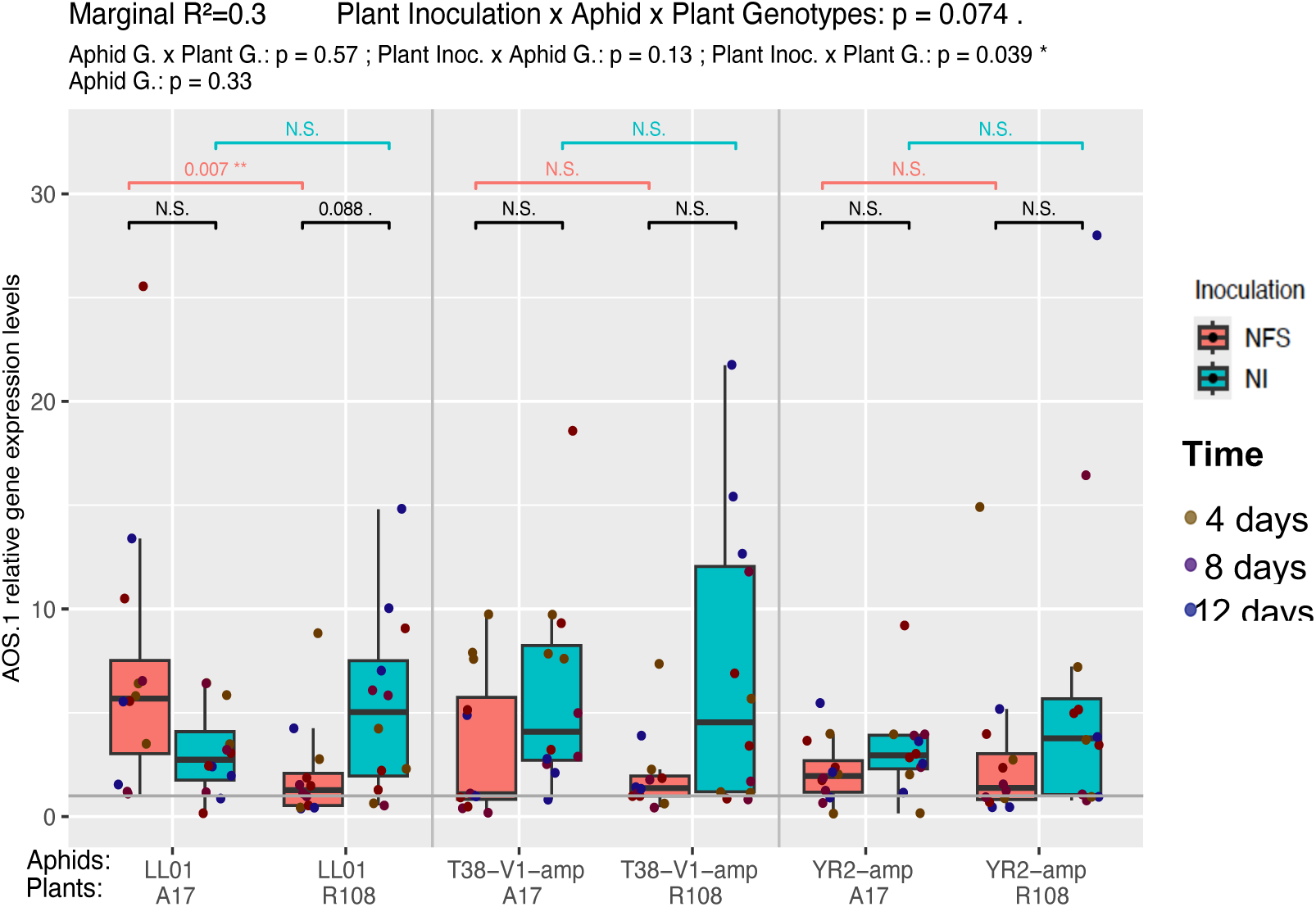
*AOS.1* gene expression in *Medicago truncatula* shoots following aphid infestation and inoculation treatment. Expression levels of the salicylic acid (SA) pathway marker gene *AOS.1* in shoots of two *Medicago truncatula* genotypes (A17 and R108) infested with three aphid genotypes (LL01, YR2-amp, T38-V1-amp), under two nitrogen treatment: nitrate-fed (NI; blue) and rhizobia-inoculated (NFS; pink) (n=3; each replicate a pool of 6 plants). Expression values are rescaled relative to non-infested control plants (grey horizontal line), which represent basal expression levels. Box plots show median and interquartile range (IQR); dots represent individual biological replicate for *AOS.1* gene expression levels observations at discrete time points post-infestation (see time colours on the legend). Mixed model predictions for the interaction of plant inoculation, aphid and plant genotype on *AOS.1* gene expression (Marginal R² = 0.3)

**Figure S6.**
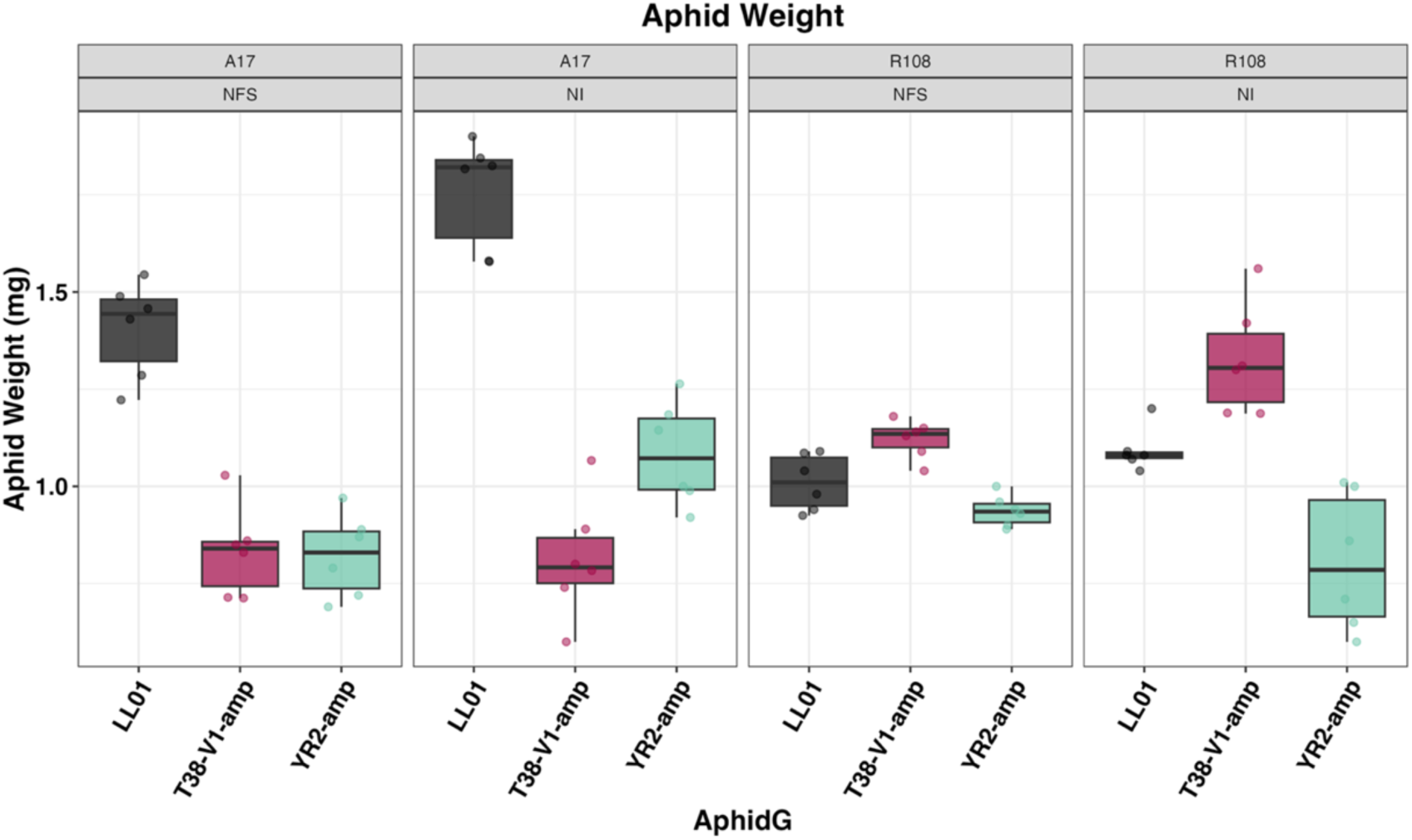
Adult aphid genotypes weight at 12 days post infestation on different *M. truncatula* genotypes under the two different nitrogen treatments. Box plots show the distribution of the mean weight in mg of living pea aphids at 12 days post infestation (*Acyrthosiphon pisum* clones LL01, YR2-amp, and T38-V1-amp) on two *Medicago truncatula* genotypes (A17 and R108) under two nitrogen treatment: mineral nitrate supplementation (NI) and rhizobial inoculation (*Sinorhizobium meliloti*; NFS). (n =6) Jittered points represent individual biological repeat for mean weight of the living aphids reached at 12 days.

**Figure S7:**
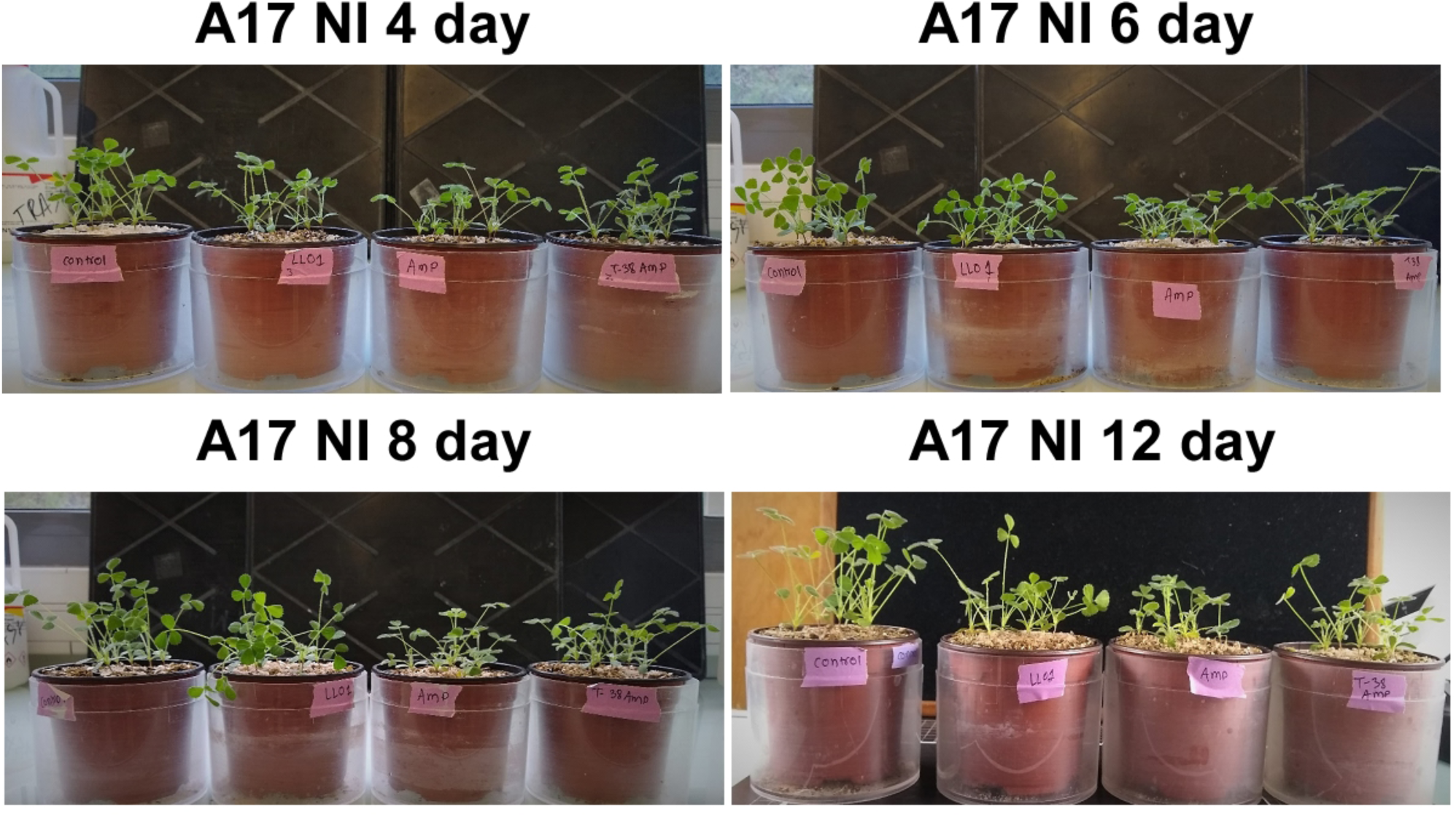
Photos of aphid-infested *M. truncatula* A17 NI plants and control plants at different time points after infestation **(A)** 4 days **(B)** 6 days **(C)** 8 days **(D)** 12 days. pots left to right: control; LL01; YR2;T38.

**Figure 8:**
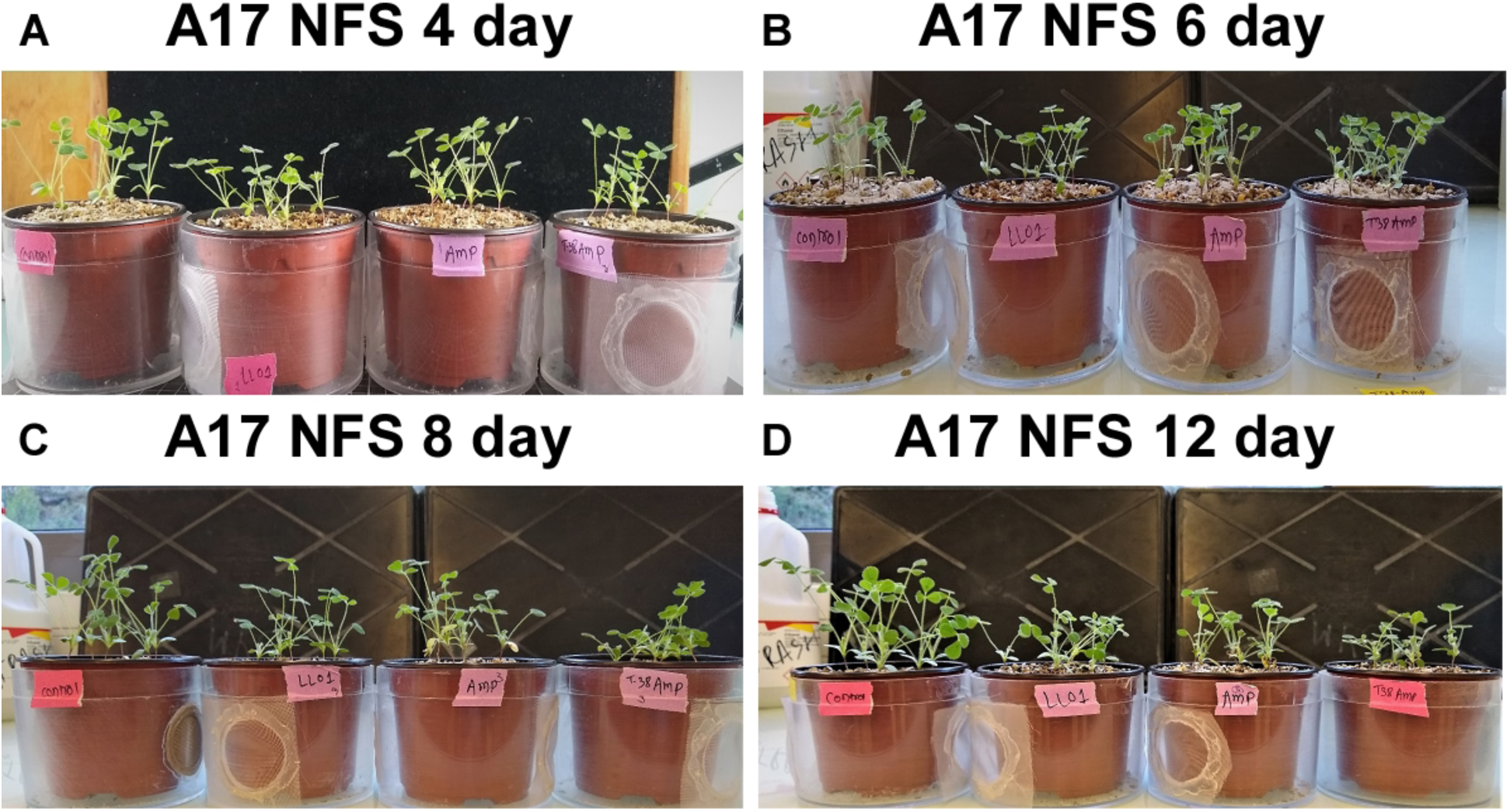
Photos of aphid-infested *M. truncatula* A17 NFS and control plants at different time points after infestation **(A)** 4 days **(B)** 6 days **(C)** 8 days **(D)** 12 days. Pots left to right: control; LL01; YR2;T38.

**Figure S9:**
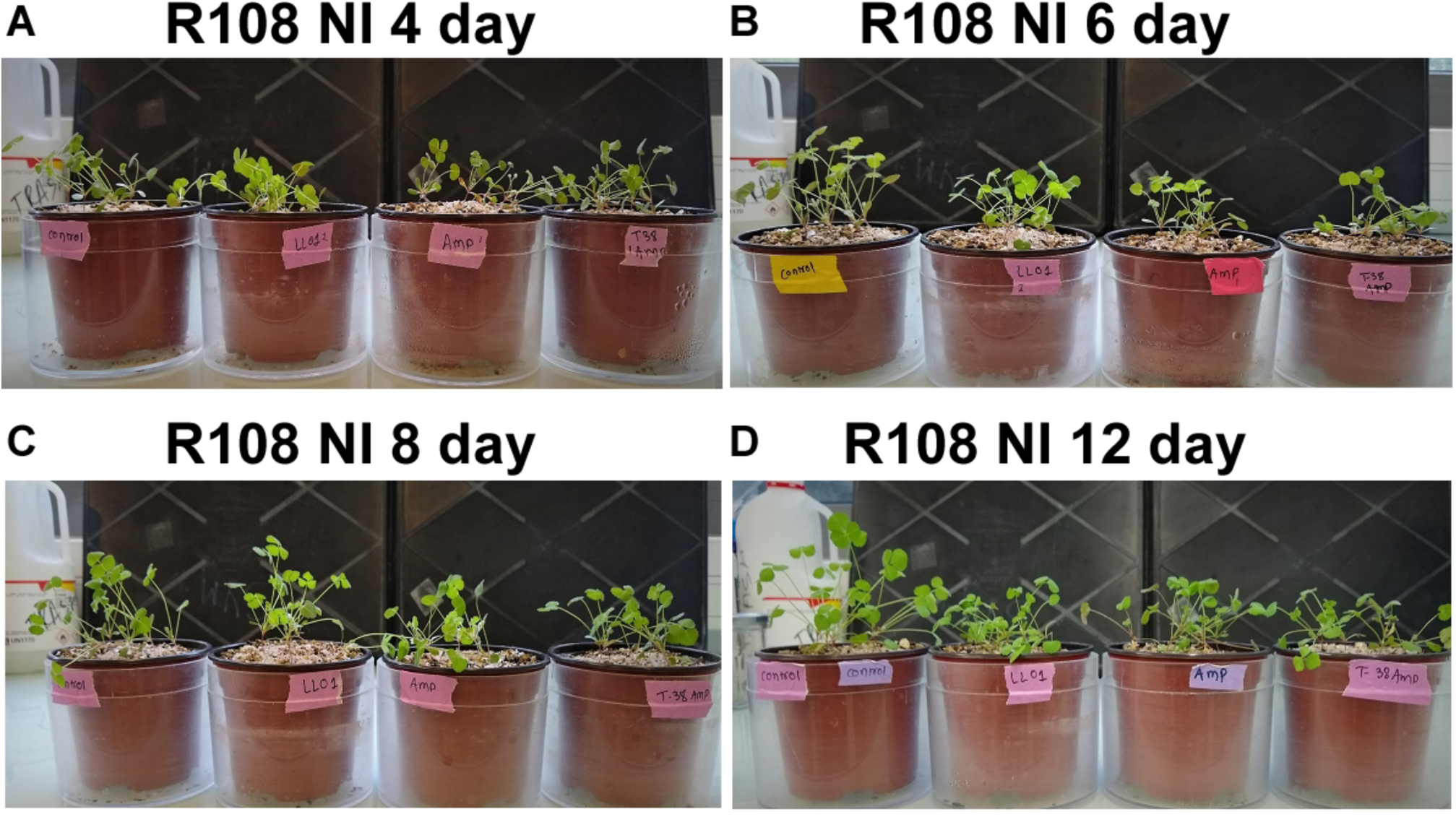
Photos of *M. truncatula* aphid-infested and control R108 NI plants at different time points after infestation. **(A)** 4 days **(B)** 6 days **(C)** 8 days **(D)** 12 days. Pots left to right: control; LL01; YR2; T38.

**Figure S10:**
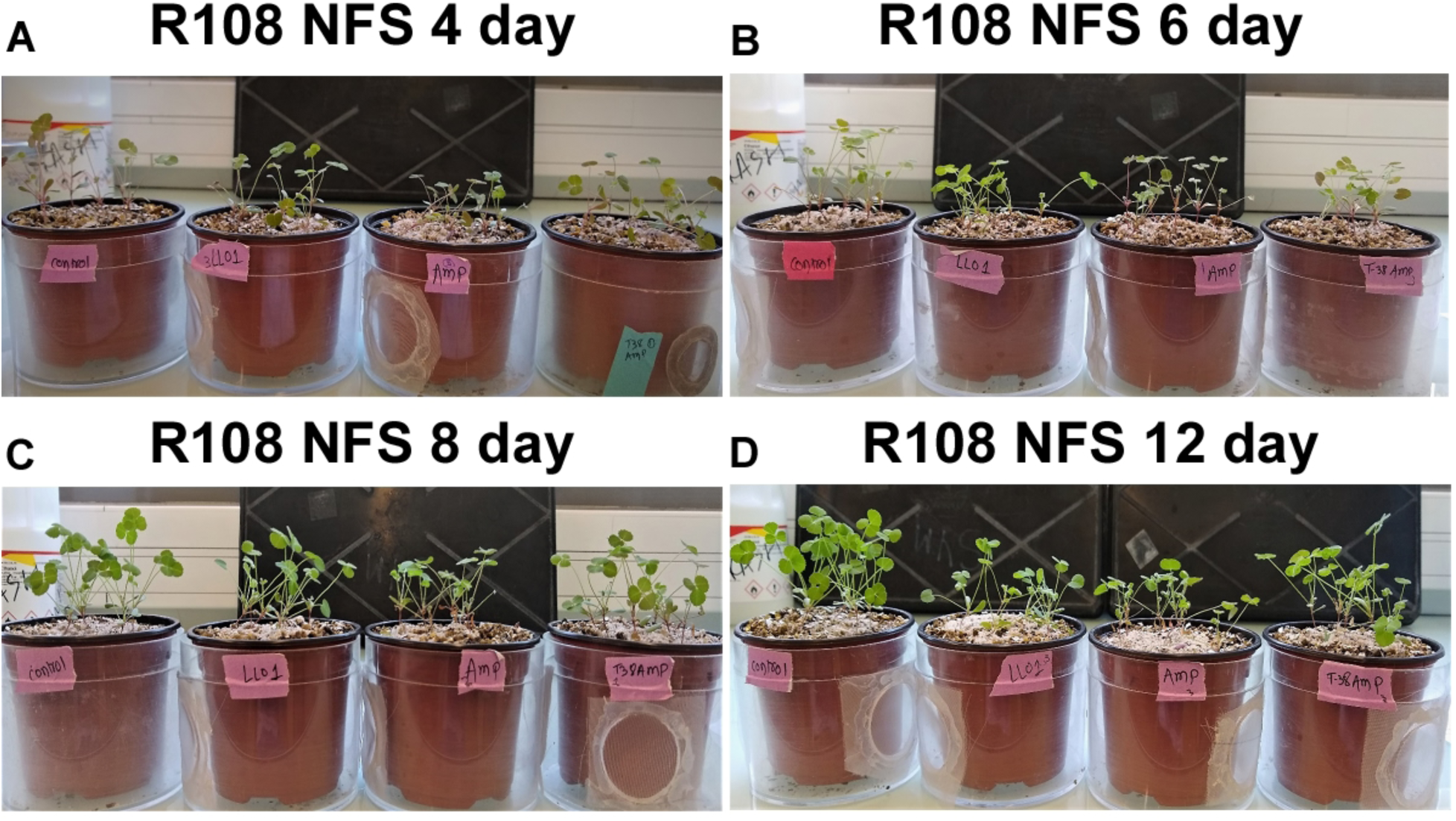
Photos of *M. truncatula* aphid-infested and control R108 NFS plants at different time points after infestation. **(A)** 4 days **(B)** 6 days **(C)** 8 days **(D)** 12 days. Pots left to right: control; LL01; YR2; T38.

**Figure S11:**
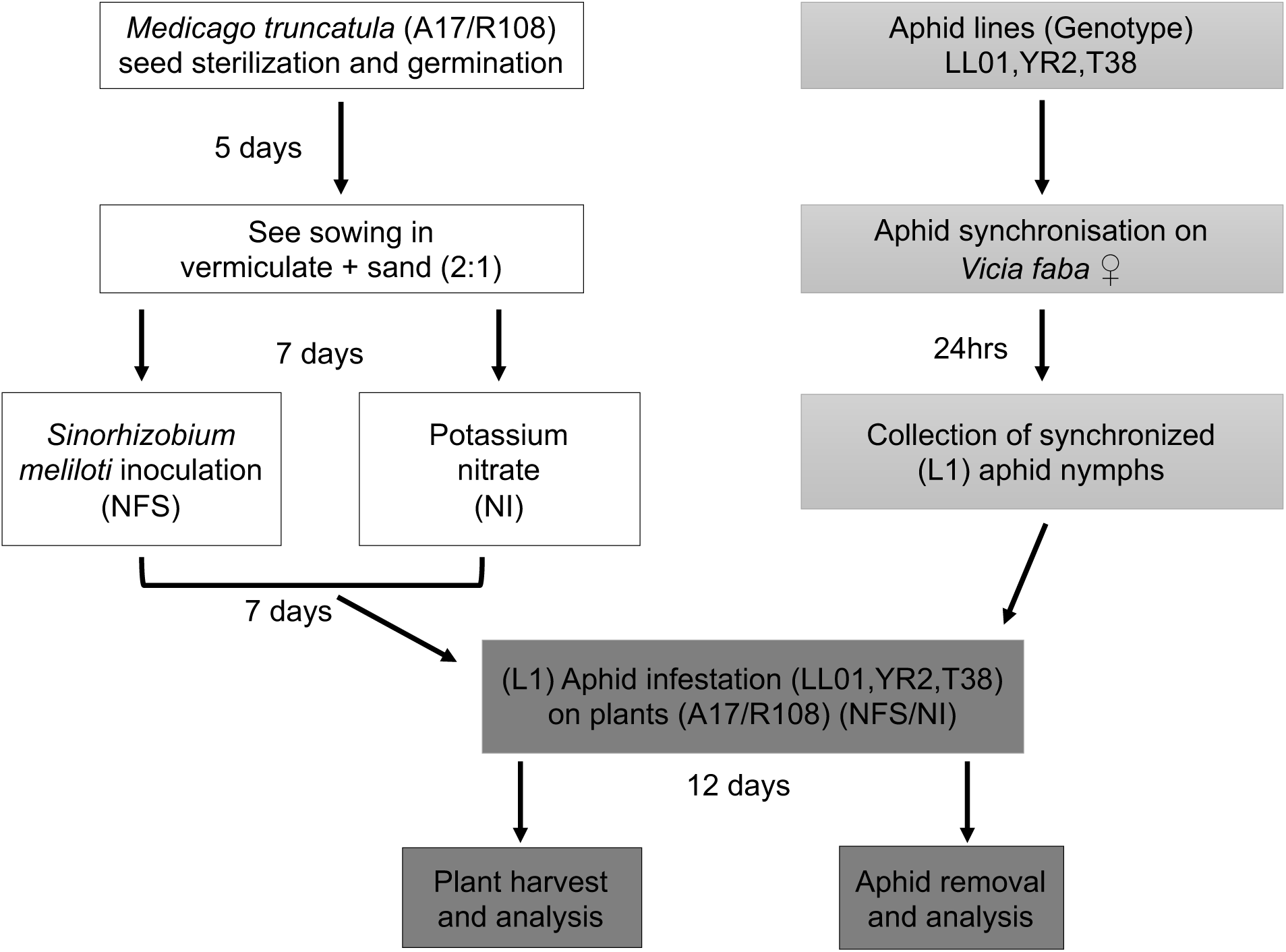
Experimental design, plants genotype and aphids genotype and inoculation time

## Supplementary Tables

**Table S1:** Origin of the aphid lines.

| Line | Color | Plant of collection | Location | Collection date | Secondary symbiont |
| --- | --- | --- | --- | --- | --- |
| LL01 | Green | Alfalfa | Lusignan, France | January 1987 | none |
| YR2-Amp* | Pink | Clover | York, UK | December 2002 | none |
| T3-8V1-Amp* | Green | Clover | Domagne, France | June 2003 | none |

\* Aphid lines were treated with ampicillin in 2010 to remove the secondary symbionts. All lines were secondary symbionts free.

**Table S2:**
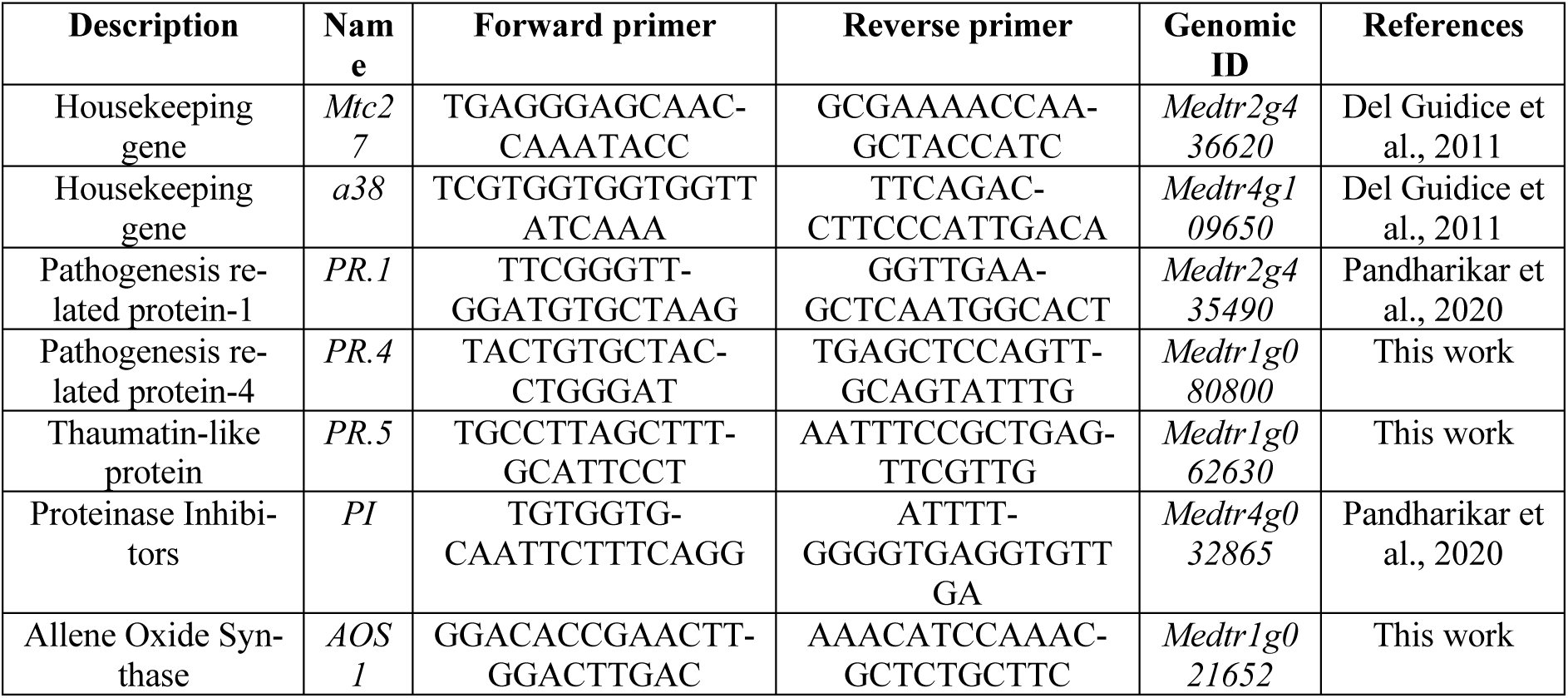
Primer sequences and conditions for qPCR analysis. Real-time qPCR was performed as follows (AriaMx Real-time PCR machine, Agilent): 95°C for 3 min followed by 40 cycles at 95°C for 5 sec and 60°C for 30 sec. The primers efficiency was evaluated on a slope of a standard curve generated using a serial dilution of the samples. Cycle threshold values (Ct) were normalized to the average Ct of two housekeeping genes *Medtr2g436620* also named *MtC27* (the homolog *of M. sativa* translationally controlled tumour protein *Msc27*), and *Medtr4g109650*.1 also named *a38* coding for a hypothetical protein. The expression of these two genes was not affected by the treatments in our experiments. The original Ct values were obtained from the machine software (Ariamix software; Agilent).

**Table S3:**
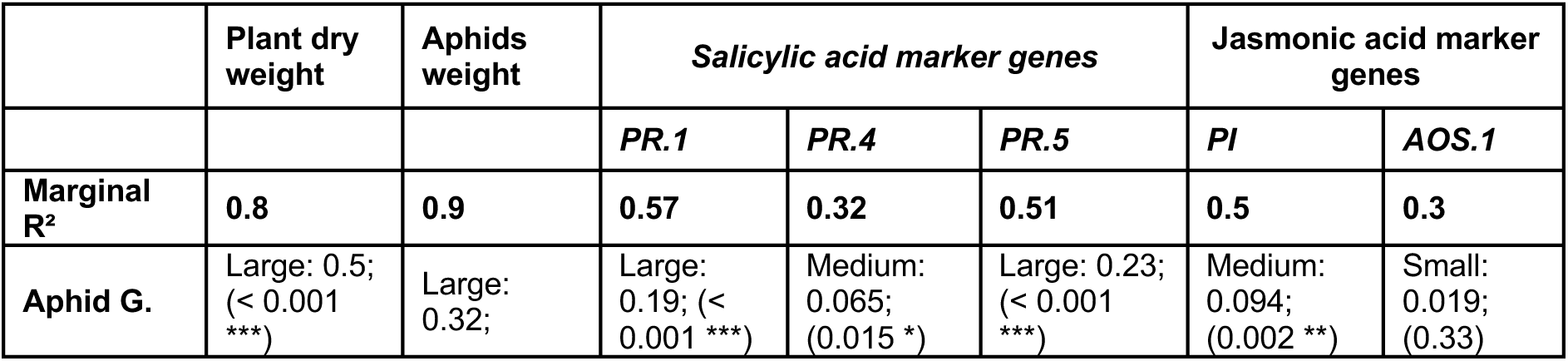
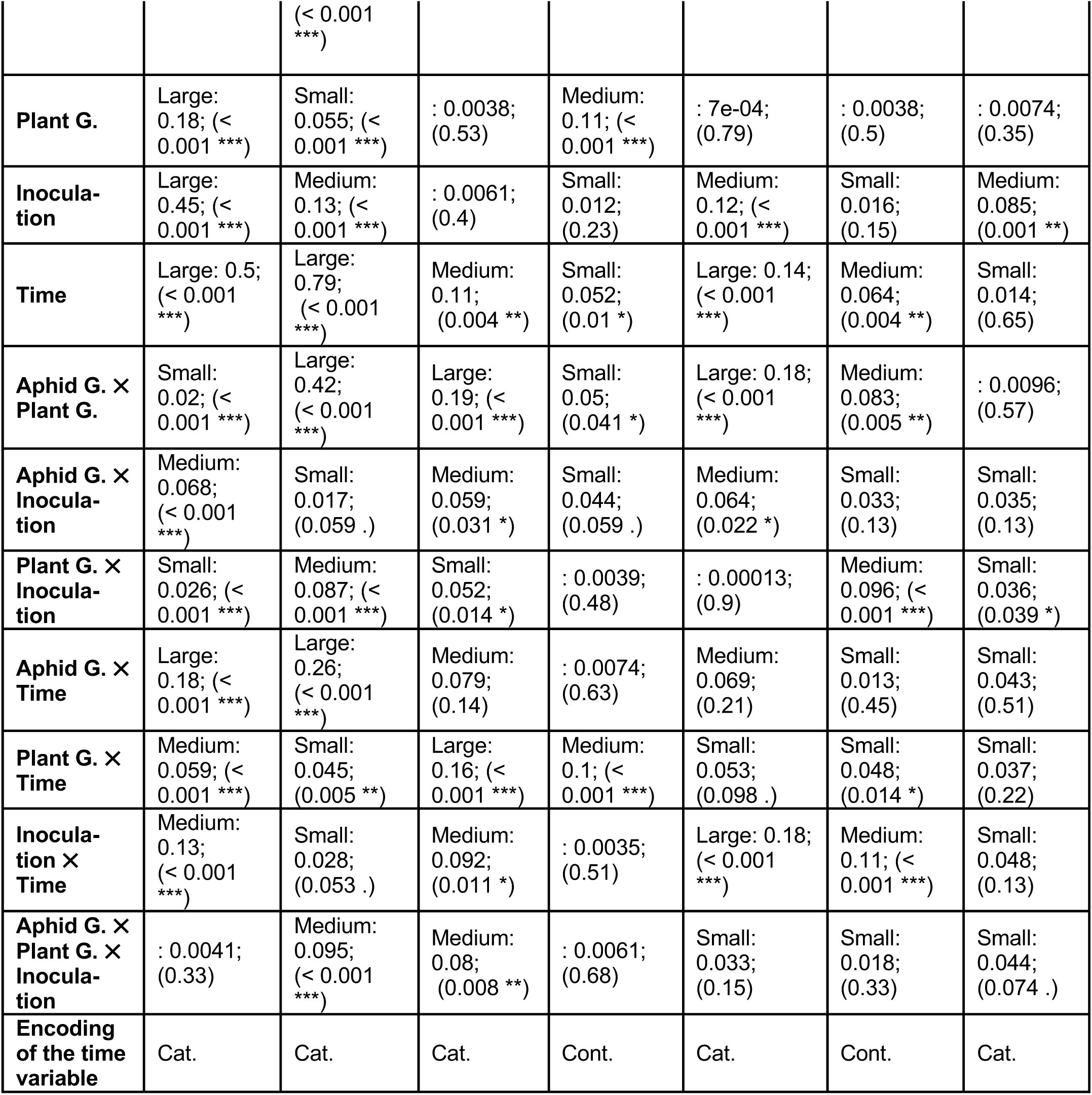
Summary of the models testing effects of the aphid and plant genotypes and of inoculation by *S. meliloti* on the plant - aphids interaction. Each column corresponds to one explained variable describing the plant - aphids interaction, and rows give the explanatory variables and their interactions, except the last row that indicate if the time was encoded as a categorical or a continuous variable. Significance levels are indicated as follows: ***p < 0.001; **p < 0.01; *p < 0.05; p < 0.1 ·p < 0.1 (marginal significance); ;ns = not significant. Effect size interpretations follow standard thresholds (η²: small ≥ 0.01, medium ≥ 0.06, large ≥ 0.14, values < 0.01 are reported numerically to indicate negligible effects).

